# Integrated Multi-analytical Framework for Comprehensive Characterization of Lignocellulosic Hydrolysates

**DOI:** 10.1101/2025.09.12.675780

**Authors:** Sarvesh Kumar, Paula Nousiainen, Donya Kamravamanesh, Rahul Mangayil

## Abstract

Lignocellulosic hydrolysates are promising second-generation feedstocks for sustainable biomanufacturing. However, biomass compositional complexities and a lack of analytical framework, particularly for lignin fraction (LF), have limited holistic characterization and impeded downstream valorization. In this study, we present an integrated multi-analytical framework for in-depth compositional profiling of spruce sawdust hydrolysate (SSH). Combining high-performance anion-exchange chromatography and liquid chromatography, sugars and inhibitors were quantified, with monomeric arabinose (78.8±0.6 g/L) and acetic acid (14.9±0.1 g/L) as predominant components, respectively. Additionally, we developed a workflow to quantify and evaluate lignin structures and interlinkages, determining factors in biomanufacturing using a unique spectroscopic-chromatographic approach enabling a comprehensive understanding of the LF. UV-Vis spectroscopy quantified a total aromatic content of 58.2±0.1g/L, while quantitative ^31^P NMR clarified the hydroxyl group distribution in the LF. ^1^H-^13^C HSQC NMR provided structural identification of lignin units and interconnecting linkages, and gel permeation chromatography showed the molecular weight distribution. Gas chromatography-mass spectrometry identified and quantified 38 lignin-derived monomers and extractives, with benzoic acid (2.7±0.2 mg/L) as the most abundant. This integrated approach provides holistic biomass compositional insights and cross-validation across methods, overcomes the limitations of a single technique, and delivers a robust workflow to evaluate the biomass suitability for bio-valorization.

**ABSTRACT GRAPHIC:** 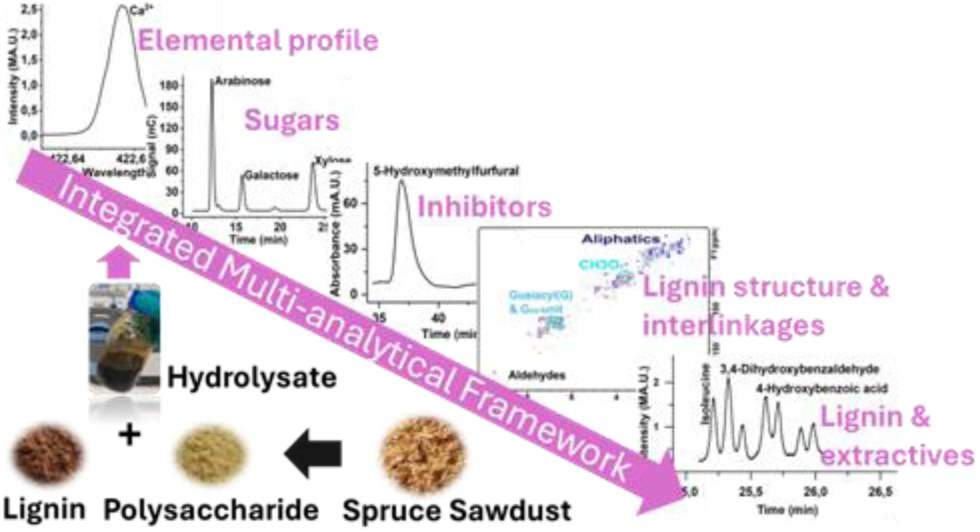

**SYNOPSIS:** This study develops an integrated multi-analytical workflow to comprehensively profile biomass hydrolysate for downstream valorization.

## INTRODUCTION

According to the *European Union Bioenergy Sustainability Report 2024*, approximately one billion cubic metres of lignocellulosic residues were generated across the 27 EU-member states, with a large portion burned for energy generation.^1^ To support sustainability objectives, there is an immense demand to develop integrated biorefinery concepts based on second-generation feedstocks, converting lignocellulosic residues into value-added bioproducts.^2^ However, the complex and recalcitrant structure of lignocellulosic residues often hinders their efficient valorization.^2,3^ To overcome this, various pretreatment methods, including acid hydrolysis, alkaline treatment, enzymatic saccharification, steam explosion, and ammonia fiber expansion, have been employed to release the lignocellulosic monomeric constituents, facilitating subsequent valorization.^4–6^

Each biomass pretreatment method has a profound impact on the treatment efficiency and hydrolysate composition. This includes fermentable sugars, microbial inhibitory compounds [furan aldehydes, organic acids, and lignin-derived monomers and oligomers (LDMOs)], and extractives that can influence downstream valorization.^2^ For instance, a comparative study by Yu et al.^7^ reports that while mild-acid hydrolysis of corn stover yielded high monomeric sugar (23.0-32.7 g/L) and microbial inhibitor (3.3-11.3 g/L; furfural, hydroxymethylfurfural, formic acid and acetic acid) concentrations, alkali and sulfite pretreatments produced lower monomeric sugars (0.8 g/L) and inhibitory compounds (3.9 g/L and 3.6 g/L, respectively).^7^ Elucidation of the compositional complexities and inhibitory compounds in biomass hydrolysates is critical for subsequent downstream bio-valorization. For example, studies focusing on biomass valorization using *Clostridia* have reported furfural 0.3 g/L, hydroxymethylfurfural 0.6 g/L^8^, and levulinic acid (LA) 0.1 g/L^9^ as potential inhibitory compounds for microbial growth and enzymatic activity. Although literature reports the capacity of acetogens to utilize the methyl groups in the methoxylated LDMOs^10^, microbial growth is inhibited at industrially relevant concentrations.^11^ The lot-to-lot variability in hydrolysate composition, together with the lack of holistic analytical frameworks, underscores the need for a quantitative workflow to assess feedstock suitability before process design for microbial valorization.

A variety of spectroscopic and chromatographic techniques have been employed to analyze lignocellulosic hydrolysate. For example, high-performance anion-exchange chromatography with pulsed amperometric detection (HPAEC-PAD) and high-performance liquid chromatography (HPLC) are the two commonly used techniques to quantify monomeric sugars and organic acids, respectively.^12–14^ CHNS/O elemental analysis and inductively coupled plasma optical emission spectrometry (ICP-OES) have been used to determine the elemental composition and inorganic salt content in the hydrolysate.^15,16^ While UV-visible spectroscopy provides an estimate of the total aromatic content, advanced analytics such as gel permeation chromatography (GPC), gas chromatography-mass spectrometry (GC-MS), Fourier transform infrared spectroscopy (FTIR), and advanced nuclear magnetic resonance (NMR) spectroscopic techniques including proton (^1^H) NMR, proton-carbon heteronuclear single quantum coherence (^1^H-^13^C HSQC) NMR and phosphorus-31 (^31^P) NMR offer in-depth compositional and structural insights of the lignin fraction (LF).^17–19^ Despite their availability and complementary capabilities, these techniques are seldom integrated into a single cohesive analytical framework, limiting comprehensive, component-level understanding of lignocellulosic hydrolysates.

The present study develops an integrated multi-analytical framework designed to address the compositional complexity of lignocellulosic hydrolysates with an aim to evaluate their suitability as feedstock for microbial valorization. By applying a suite of complementary chromatographic and spectroscopic techniques, the resulting integrative approach offers a holistic and quantitative understanding of hydrolysate composition. The overarching goal of this study was to establish an integrative workflow and analytical benchmarks to assess lignocellulosic hydrolysate components, specifically the LF, thereby supporting the development of sustainable bioprocessing strategies. To demonstrate the applicability of this methodology, we present a case study using spruce sawdust hydrolysate (SSH) derived from pressurized hot water extraction of spruce sawdust, while maintaining broader relevance for diverse lignocellulosic feedstocks.

## MATERIALS AND METHODS

### Biomass and Chemicals

The lignocellulosic hydrolysate, referred to as spruce sawdust hydrolysate (SSH), was the liquid fraction obtained after pressurized hot water extraction (PHWE) of pretreated spruce sawdust at 160 °C for 20 min, following separation of the polysaccharide and polyphenolic lignin products. SSH was provided in a concentrated form by Boreal Bioproducts (Espoo, Finland). For analysis of SSH, the chemicals used, the extracted soluble lignin fraction (LF) from the SSH and the sample preparation procedures for analytical measurements are provided in the Supporting Information (Supporting materials and methods). All analyses were performed in triplicate unless otherwise noted. ^1^H-^13^C HSQC NMR (semi-quantitative), FTIR, and ^1^H NMR spectra were acquired from a single sample; multiple scans were averaged to improve spectral quality.

### Analyses of SSH Components

#### Elemental Composition

Elemental analysis was conducted using a Flash Smart CHNS/O Elemental Analyzer (Thermo Fisher Scientific, USA) following the protocol described by Robertson et al.^20^ Freeze-dried SSH (2 mg) was analyzed with sulfanilamide as a standard for CHNS quantification, and 2,5-bis(5-tert-butyl-2-benzoxazolyl) thiophene for oxygen quantification using helium as the carrier gas.

#### Inorganic Salts

ICP-OES (5900 SVDV, Agilent, USA) was used to quantify the inorganic salts (Ca^2+^, Cu^2+^, Fe^3+^, K^+^, Mn^2+^, Na^+^, Pb^2+^, and Zn^2+^) present in SSH. Instrument calibration and quality control standards were conducted using Sc^+^ and Se^+^ to ensure the accuracy and precision of the measurements.

#### Monomeric Sugars

HPAEC-PAD (ICS-5000, Thermo Fisher Scientific, USA) containing a gold working electrode and an Ag/AgCl reference electrode, CarboPac PA20 analytical column (3 × 150 mm, Thermo Fisher Scientific, USA) and Milli-Q water as the mobile phase (flow rate, 0.37 mL/min) was used to analyze the monomeric sugars in the SSH fraction using the method described by Rohrer et al.^21^ The column washing and equilibration between the runs were performed using 0.2 M NaOH and Milli-Q water, respectively.

#### Organic Acids, Furans and Aromatics

Acetic acid, formic acid, furfural, hydroxymethyl furfural (HMF), and levulinic acid were analyzed using HPLC (Ultimate 3000, ThermoFisher Scientific, USA) and established protocols described by Wrigstedt et al.^22^ The HPLC was equipped with UV and refractive index detectors (RID) and a Rezex ROA-Organic Acid H^+^ (8%, 300 x 7.8 mm, Phenomenex, USA) column. The column oven and RID temperature were set at 55 °C. UV detection was performed at 210 nm, 254 nm, and 272 nm. The mobile phase used was 5 mM H_2_SO_4_ at a flow rate of 0.5 mL/min.

Total aromatic content in SSH was assessed using a UV-Visible spectrophotometer (UV-2550, Shimadzu, Japan) by measuring UV absorbance at 205 nm with a known extinction coefficient for spruce^23^ based on the principles of the TAPPI UM 250.

#### Lignin derived Monomers, Oligomers, and Extractives

^1^H NMR and ^1^H-^13^C HSQC NMR analyses were performed to characterize lignin-derived monomers and oligomers (LDMOs) interunit linkages and structural distribution in the LF. NMR analysis was conducted using a Bruker AV III 400 MHz NMR Spectrometer (Bruker, Germany) with a 5 mm BBFO iProbe for spectra collection. For ^1^H NMR analysis, a standard pulse sequence of zg30 with 16 scans was used, with a pulse delay of 4 seconds and a spectral width of 16 ppm. For ^1^H-^13^C HSQC NMR analysis, a hsqcetgpsisp.2 standard pulse sequence was used with a 2-second pulse delay, and 24 scans were collected with spectral widths of 11 ppm in F2 (^1^H) dimension with 1024 data points, and 215 ppm in F1 (^13^C) dimension with 128 data points. The spectra were referenced to the residual dimethyl sulfoxide DMSO solvent signal at δC/δH 39.52/2.50 ppm. All spectra were analysed with Bruker Topspin 4.4.0 MAC version using standard processing parameters.

To quantify the aliphatic and aromatic hydroxyl groups in the LF, ^31^P NMR analysis was conducted. The zgig pulse program was done at 25 °C, with parameters set to 128 scans, 1-second acquisition time, 5-second pulse delay, 90° pulse angle, and a total acquisition time of 13 minutes. Data was processed using a line broadening factor of 5 Hz.

To identify the functional groups and overall chemical bonding patterns of LDMOs in the LF, the vacuum-dried extraction of the LF was analyzed using a Spectrum Two FTIR spectrometer (PerkinElmer, USA) equipped with an attenuated total reflection (ATR) module. Briefly, vacuum-dried LF sample was placed on a diamond crystal plate, and a standard high-performance Lithium tentalate (LiTaO3) mid-infrared (MIR) detector was used to measure the spectrum range of 4000 to 450 cm^-1^ with a resolution of 4 cm^-1^. For data collection, a standard optical system with a KBr window was used; spectra were recorded as an average of 30 scan accumulations per sample.

Quantification of lignin-derived monomers and extractives in the LF was conducted using a GC-MS (QP2010SE, Shimadzu, Japan) equipped with an HP-5 (30m x 0.25 mm x 0.25 µm, Agilent, USA) nonpolar column and helium as the carrier gas (flow rate, 1 mL/min). Briefly, samples were injected in a splitless mode, and the injector temperature was 300 °C. GC-MS analysis was performed using the following oven temperature program: an initial temperature of 80 °C held for 5 minutes, followed by a ramp of 4 °C/min to a final temperature of 300 °C. The final temperature was held for 20 minutes, resulting in a total run time of 80 minutes. GCMSsolution and LabSolutions software with the NIST 2017 (National Institute of Standards and Technology), and GC-MS mass spectral library were used for data analysis. Column cleaning using dichloromethane washing and pyridine injection was conducted between the analysis runs. A set of 20 lignin-derived monomers standards was used for analysis, with 4-hydroxyacetophenone and veratric acid as internal standards (0.5 mg/mL), as indicated in the Supporting Information file (Supporting materials and methods). Final concentrations of lignin-derived monomers and extractives were calculated and expressed as total concentrations in SSH.

To analyze the molecular weight distribution of the LDMOs in LF, GPC was performed using an Agilent 1100 HPLC system (Agilent, USA) equipped with two Phenomenex Phenogel SEC columns (300 mm×7.8 mm, 5 μm particle size, 5 nm and 100 nm pore sizes, Phenomenex, USA) and a UV detector set at 280 nm for compound detection. Tetrahydrofuran was used as the mobile phase at a constant flow rate of 1.0 mL/min. The analyses were conducted under ambient conditions.

#### Linearity of Detection

The relationship between the acquired signal (peak area) and analyte concentration obtained from the tested analytics was evaluated using linear regression, with an acceptable linearity of R^2^ > 0.99 (regression coefficient).

## RESULTS AND DISCUSSION

The primary objective of this study was to develop a comprehensive analytical framework for in-depth profiling of lignocellulosic hydrolysates. SSH was selected as a representative biomass sample to demonstrate the importance of systematic characterization of key hydrolysate components and their relevance in evaluating hydrolysate suitability for downstream valorization.

### Total Solids and Elemental Composition of the Hydrolysate

The ICP-OES analysis of SSH revealed a diverse range of metal ions. Ca^2+^ exhibited the highest concentration of 835.9±160.7 mg/L, followed by notable amounts of K^+^, Fe^3+^, Mn^2+^, Na^+^, Zn^2+^, and trace levels of Pb^2+^ and Cu^2+^ **(Table 1)**. The prevalence of these metal ions could either be attributed to their abundance in the biomass or to their introduction during the pretreatment process.

**Table 1.** Concentration of Metal Ions and Elemental Composition in SSH.

| Entities | Metal Ions | Concentration (mg/L) |
| --- | --- | --- |
| Inorganic salts | Ca<sup>2+</sup> | 835.9±160.7 |
|  | Cu<sup>2+</sup> | 0.01±0.04 |
|  | Fe<sup>3+</sup> | 31.1±0.5 |
|  | K<sup>+</sup> | 273.5±22.2 |
|  | Mn<sup>+</sup> | 66.6±0.2 |
|  | Na<sup>+</sup> | 14.9±0.2 |

|  | Pb<sup>2+</sup> | 0.2±0.4 |
| --- | --- | --- |
|  | Zn<sup>2+</sup> | 9.2±0.1 |
|  | Elements | Concentration (% w/w) |
| Elemental composition | C | 40.5±0.3 |
|  | H | 6.2±0.02 |
|  | N | 0.2±0.01 |
|  | S | 0.02±0.01 |
|  | O | 48.7±0.4 |

In contrast to the conventionally used flame atomic emission spectroscopy (AES), ICP-OES facilitates the simultaneous quantification of both major and trace metal ions in a single analytical run, without the need for pre-concentration or extensive sample cleanup.^24^ Furthermore, ICP-OES outperforms AES in sensitivity and spectral interferences, and performs robustly in organic-rich and complex matrices such as lignocellulosic hydrolysates^25^, and thus was selected as the method of choice in this study. Metal-ion availability is a key determinant of microbial growth, product formation, and inhibition. This is particularly critical for anaerobic microbial cell factories such as *Clostridium carboxidivorans*^26^, and *C. ljungdahlii*^27^, in which metalloenzymes requiring specific trace-metal cofactors underpin central metabolism and product formation.^27^ For instance, in *C. carboxidivorans*, Ni^2+^ and Fe^2+^ promote microbial growth, whereas molybdate inhibits both growth and restricts the carbon flow through the ethanol production pathway.^26^ In the solventogenic bacterium *C. acetobutylicum*, a well-known biofuel producer, supplementation with calcium salts in hydrolysate increases intracellular Ca^2+^ levels and enhances acetone-butanol-ethanol (ABE) production.^28^ Given that hydrolysates inherently contain various metal ions, a comprehensive analysis of their concentrations is essential to optimize hydrolysate-based microbial valorization processes.

To complement the ionic profile, the total solid-phase elemental composition was assessed using CHNS/O elemental analysis. The analysis revealed that SSH contained high concentrations of carbon and oxygen, followed by lower concentrations of hydrogen, and trace amounts of nitrogen and sulfur **(Table 1)**, indicating SSH contains organic matter essential for microbial fermentation. These values are consistent with the elemental compositions of spruce biomass as reported in literature, which typically range from 40-42% carbon, 5.5-6.5% hydrogen, <0.2% nitrogen, and trace levels of sulfur.^29^ Importantly, elemental profiling of biomass substrates is critical in the precise calculation of carbon mass balance and yields in bioconversion studies, ensuring that the performance metrics of microbial strains or bioprocess workflows are based on a complete and chemically validated substrate profile.

### Compositional Analysis of Sugars and Microbial Inhibitors Using Chromatographic Techniques

A mixture of monomeric and oligomeric sugars in SSH was detected by HPAEC-PAD, revealing an overall carbohydrate profile, including arabinose, rhamnose, galactose, glucose, xylose, and mannose. Monomeric sugars accounted for approximately 29% of total sugars, with an arabinose content of 78.8±0.6 g/L constituting ∼50% of the total monomeric sugar content **(Table 2)**. Other monomeric sugars, such as glucose, galactose, xylose, mannose, and rhamnose, were also detected, with xylose (38.1±0.3 g/L) being the second most abundant **(Table 2)**. To determine the total sugar content, an additional acid hydrolysis step using the NREL laboratory analytical procedure was performed (Supporting Information, Materials and Methods, Sample Preparation). The quantity of oligomeric sugars was determined by subtracting the concentration of monomeric sugars from the quantified total sugars and expressed in terms of the concentration of the corresponding monomeric sugars. The quantitative profiling of oligomeric sugars revealed an abundance of oligomeric mannose and xylose in SSH. The detection of mannose (134.0±7.2 g/L), glucose (32.5±1.5 g/L), and galactose (45.7±2.5 g/L) in the hydrolyzed biomass in oligomeric form indicates the presence of mannan and glucomannan oligosaccharides from acetylated galactoglucomannan hemicellulose.^30,31^ Additionally, substantial concentrations of oligomeric xylose (133.6±7.5 g/L), arabinose (42.6±4.8 g/L), and rhamnose (2.2±0.3 g/L) suggest the presence of xylo-oligosaccharides, arabinoxylooligosaccharides, and pectins in the biomass.^32^ Analysis of oligosaccharides alongside monomeric sugars is essential but often overlooked, as diverse microbes such as *Bifidobacterium* and *Lactobacillaceae* can metabolize such substrates to yield value-added products.^33,34^

**Table 2.**
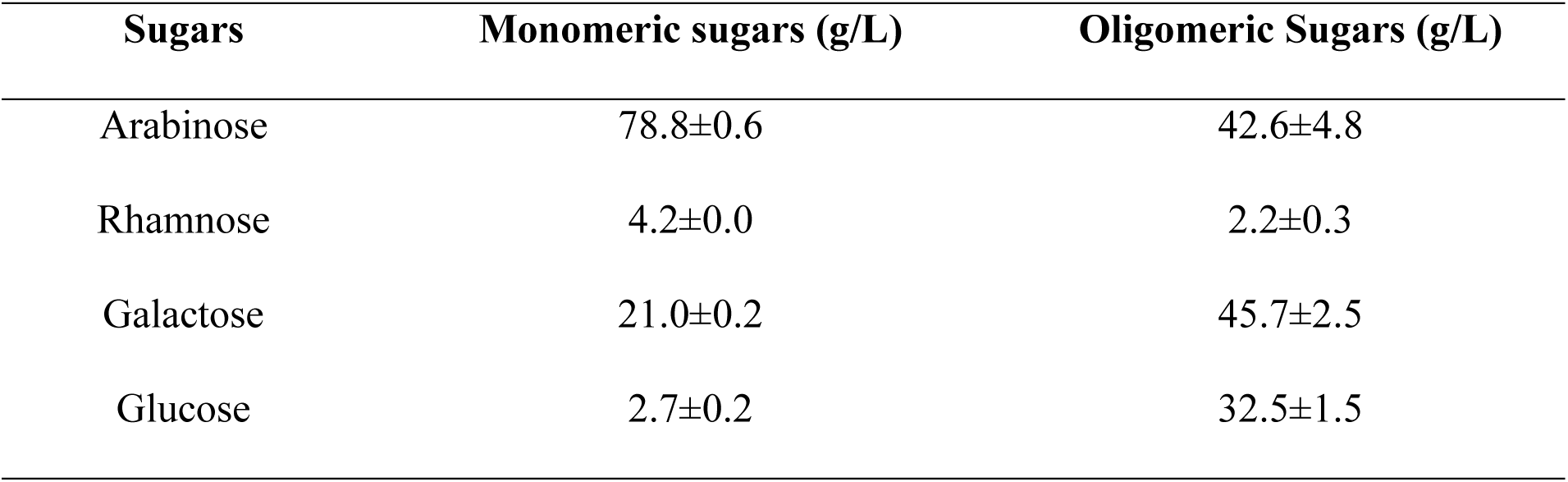
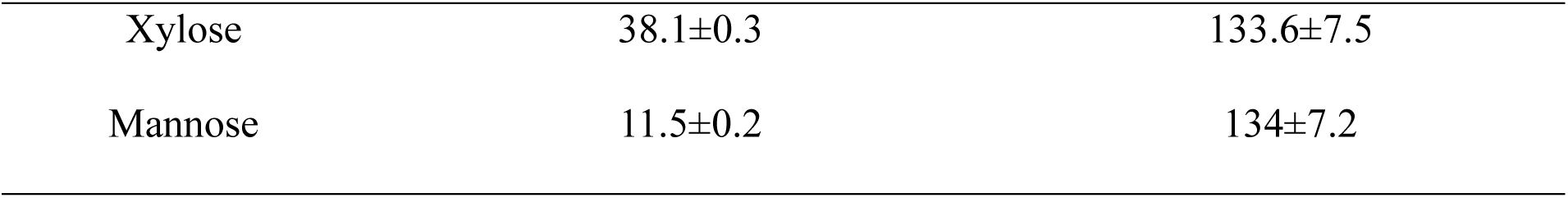
Monomeric and Oligomeric Sugar Concentrations in SSH.

| Sugars | Monomeric sugars (g/L) | Oligomeric Sugars (g/L) |
| --- | --- | --- |
| Arabinose | $78.8 \pm 0.6$ | $42.6 \pm 4.8$ |
| Rhamnose | $4.2 \pm 0.0$ | $2.2 \pm 0.3$ |
| Galactose | $21.0 \pm 0.2$ | $45.7 \pm 2.5$ |
| Glucose | $2.7 \pm 0.2$ | $32.5 \pm 1.5$ |
| Xylose | 38.1±0.3 | 133.6±7.5 |
| Mannose | 11.5±0.2 | 134±7.2 |

Comparative data in **Table 3** demonstrate that the sugar profile observed in this study aligns with Zhang et al.^35^, wherein the authors reported mannose as the predominant monosaccharide in spruce sawdust hydrolysates following PHWE. In contrast, Frankó et al.^36^ observed a glucose-rich composition in Norway spruce wood and bark hydrolysates, likely due to harsher pretreatment conditions (210 °C) or intrinsic differences in biomass composition. Similarly, Vallejos et al.^37^ reported that PHWE of sugarcane bagasse yielded hydrolysates containing 0.7% arabinose and 0.2% xylose (based on oven-dried biomass), highlighting the influence of biomass type and pretreatment on sugar composition. Although studies, such as those by Gunes et al.^38^ and Göncü et al.^39^, have measured total fermentable sugars spectrophotometrically, albeit without resolving individual monomeric sugars. As microorganisms display specific substrate preferences and co-utilization behaviors toward individual sugars present in complex mixtures^40^, detailed compositional information is critical for selecting suitable cell factories to optimize the biomanufacturing process.

**Table 3.** Comparison of hydrolysate composition from various lignocellulosic feedstocks and hydrolysis methods from recent studies.

| **Hydrolysate** |  | **Spruce**<br>**sawdust** | **Sugarcane**<br>**bagasse** | **Norway**<br>**spruce wood** | **Norway**<br>**spruce bark** | **Wheat**<br>**straw** | **Switch**<br>**grass** | **Pistachio**<br>**shells** | **Spruce**<br>**sawdust** |
| --- | --- | --- | --- | --- | --- | --- | --- | --- | --- |
| Hydrolysis method<br>(condition) |  | PHWE<sup>α</sup> | PHWE (160<br>°C, 20 min) | SP (210 °C, 5<br>min) | SP (210 °C, 5<br>min) | AH (190<br>°C, 10 min) | HW<sup>β</sup><br>/EH | US/EH | PHWE (160<br>°C, 10 min) |
| Analysis methods<br>(sugars & inhibitors) |  | GC-methanolysis | HPLC | HPLC | HPLC | HPLC | Spec | Spec | HPAEC-<br>PADHPLC |
| **Hydrolysate profile** |  | **Weight %** | **% ODB** | **Concentrations (g/L)** |  |  |  |  |  |
| Total<br>sugars | Arabinose | n.a. | 0.7 | 2.7 | 10 | 3.9±0.1 | 355.72<sup>δ</sup> | 7.34<sup>δ</sup> | 121.2±4.7 |
|  | Rhamnose | n.a. | n.a. | n.a. | n.a. | n.a. |  |  | 6.4±0.3 |
|  | Galactose | n.a. | n.a. | 5.1 | 4.3 | 3.1±0.6 |  |  | 66.6±2.5 |
|  | Glucose | n.a. | 0.0 | 26.5 | 18.2 | 84.9±0.8 |  |  | 35.1±1.4 |
|  | Xylose | n.a. | 0.2 | 9.7 | 6 | 32.5±0.7 |  |  | 171.6±7.4 |
|  | Mannose | 39.5-50.7 | n.a. | 21.7 | 5.6 | 1.6±0.4 |  |  | 145.4±7.2 |
| Inhibitors | Acetic Acid | n.a. | 0.4 | 1.9 | 0.6 | 6.0±0.6 | n.a. | n.a. | 14.9±0.1 |
|  | Formic Acid | n.a. | 0.1 | 1.5 | 0.4 | n.a. | n.a. | n.a. | 0.8±0.1 |
|  | Levulinic acid | n.a. | n.a. | 1 | 0.02 | n.a. | n.a. | n.a. | 1.5±0.1 |
|  | Furfurals | n.a. | 0.04 | 1.3 | 0.4 | 0.8±0.1 | n.a. | n.a. | 0.1±0.0 |
|  | 5-Hydroxy Methyl furfural | n.a. | 0.0 | 2.0 | 0.4 | 0.5±0.1 | n.a. | n.a. | 1.0±0.0 |
| Total aromatics | LDMOs and extractives | n.a. | n.a. | n.a. | n.a. | n.a. | n.a. | n.a. | 58.2±0.1<sup>⊙</sup> |
| Salts | Metal ions | n.a. | n.a. | n.a. | n.a. | n.a. | n.a. | n.a. | 8<sup>ψ</sup> |
| References |  | Zhang et al<sup>35</sup> | Vallejos et al.<sup>37</sup> | Franko et al.<sup>36</sup> | Franko et al.<sup>36</sup> | Persson et al.<sup>45</sup> | Gunes et al.<sup>38</sup> | Goncu et al.<sup>39</sup> | This study |

The HPAEC-PAD system used in the present work facilitated the precise quantification of SSH sugars with high linearity (R^2^ > 0.99) **(Table 3** **and Table S1)**. Compared to the conventional HPLC systems with UV or RI detection, HPAEC-PAD offers superior resolution of structurally similar sugars without the need for derivatization.^14,41^ While HPLC-MS enables sensitive sugar detection and tolerates some co-elution of analytes, it often necessitates complex sample preparation, isotopically labeled internal standards, and may face challenges in distinguishing SP - steam pretreatment; AH - acid hydrolysis; HE - hot water extraction; EH- enzyme hydrolysis; US - ultrasonication; lignin-derived monomers and oligomers (LDMOs); Spec - spectrophotometry; ODB - oven dry biomass α - hydrolysis at 120 and 160 °C for 20 and 60 min; β - 122.12 °C for 50.08 min; δ - Total fermentable sugars; Θ - total aromatics concentration estimated by UV-Visible spectrophotometer; ψ - metal ions **(Table 1)** analyzed by ICP-OES are presented as g/L; n.a. - not given. isobaric sugars without additional chromatographic separation.^41^ In this regard, HPAEC-PAD provides a practical approach for monitoring individual sugar consumption during microbial fermentation, offering critical insights into the impact of hydrolysate composition on microbial metabolism, an aspect often underreported in hydrolysate evaluation studies.^42–44^

### Evaluation of Organic Acids and Furans

Acetic acid, formic acid, and levulinic acid were detected at 210 nm, while furfural and HMF were detected both at 254 nm and 272 nm, respectively during analysis of microbial inhibitors with HPLC. The method showed strong linearity (R^2^ > 0.99; **Table S1),** yielding concentrations of 1.5±0.1 g/L, 0.8±.0 g/L, 1.5±0.1 g/L, 0.1±0.0 g/L, and 1.0±0.0 g/L, respectively, consistent with previous reports **(Table 3).** SSH contained low levels of microbial inhibitors (<2 g/L), indicating that it would not negatively affect the growth or fermentation capacities of microbial cell factories.^8,9^ HPLC equipped with reversed-phase long alkyl chain bonded to silica stationary phase C30 columns and UV detection has been conventionally used to analyze the inhibitory furans (furfural and HMF) in lignocellulosic hydrolysates.^46,47^ Nevertheless, coupling an H^+^ ion exclusion column with UV detection, as in this study, broadens the analytical capabilities, facilitating simultaneous detection of organic acids, alcohols, and furans inhibitors. This configuration offers versatility for real-time monitoring in microbial fermentation studies. Thus, by integrating HPAEC-PAD for sugar profiling with HPLC for inhibitor detection, a dynamic profile of biomass bioconversion can be achieved.

### Analysis of Lignin-derived Monomers, Derivatives, and Extractives Using Spectroscopic and Chromatographic Techniques

The combined application of chromatographic and spectroscopic techniques enabled a detailed and multidimensional analysis of the lignin fraction (LF) within the SSH. Each method provided complementary insights, such as molecular weight distribution, functional group composition, and structural features, which are typically obtained in isolation and rarely integrated in a single study. The integrated approach presented in this study offers a comprehensive understanding of lignin composition and structure, providing a robust foundation to evaluate its impact on downstream microbial processes and valorization strategies.

### Structural Profiling of Lignin-derived Monomers and Oligomers via Spectroscopic Analysis

Initial insights obtained using ^1^H NMR spectroscopy revealed the presence of carboxylic acids, aldehydes, phenolic hydroxyls, aromatics, oxygenated aliphatic groups, and aliphatic groups in the LF **(Figure S1)**. However, detailed structural identification was limited due to extensive signal overlap. To resolve the overlapping signals and gain deeper insights, ^1^H-^13^C HSQC NMR spectroscopy was performed. This method enabled visualization of the interconnecting side-chain linkages of LDMOs by their proton and carbon correlation signals **(Figure 1)**. Additionally, this approach facilitated semi-quantitative determination relative to the aromatic guaiacyl (G) units, which are predominant in softwood lignin. This allowed us to correlate each type of linkage based on their relative abundance expressed as percentage in relation to 100 G-units in the sample.^20^ ^1^H-^13^C HSQC analysis revealed that the LF was composed primarily of guaiacyl-derived units (79.3%) and α-oxidized guaiacyl-derived units (20.7%), the latter characterized by carbonyl functionalities at the benzylic position (e.g., aldehydes, ketones, carboxylic acids, and esters) **(Table 4)**. Among the oxygenated derivatives, aromatic aldehydes and furfurals were detected in significant amounts based on their well-isolated correlation signals of -CH=O groups with 8.2% and 3.5%, respectively, supporting the observed distribution of carbonyl-containing functionalities.

**Figure 1.**
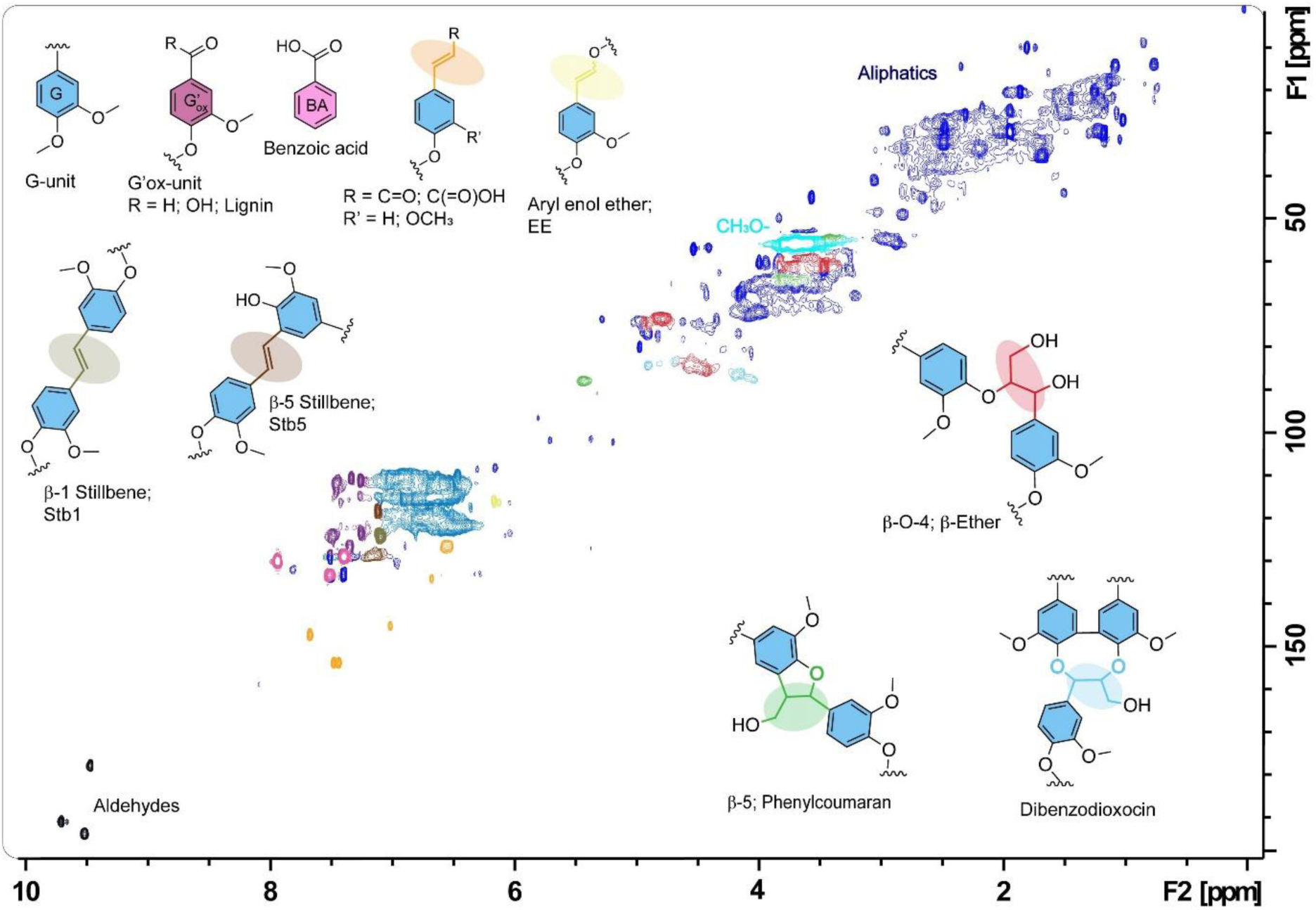
^1^H-^13^C HSQC NMR spectrum of the LF extracted from the SSH. The presented data show detailed assignments of aromatic carbons, double-bonded carbons, benzoic acid (BA), and methoxy (CH_3_O-) groups, as well as structural motifs such as guaiacyl units (G-unit), oxidized guaiacyl units (G’_ox_), and β-O-4, β-5, and dibenzodioxocin interunit linkages, as highlighted in different colors. Additional non-defined spectral regions corresponding to carbohydrates, terpenes, and aliphatic fatty acid-derived structures are indicated in dark blue.

**Table 4.**
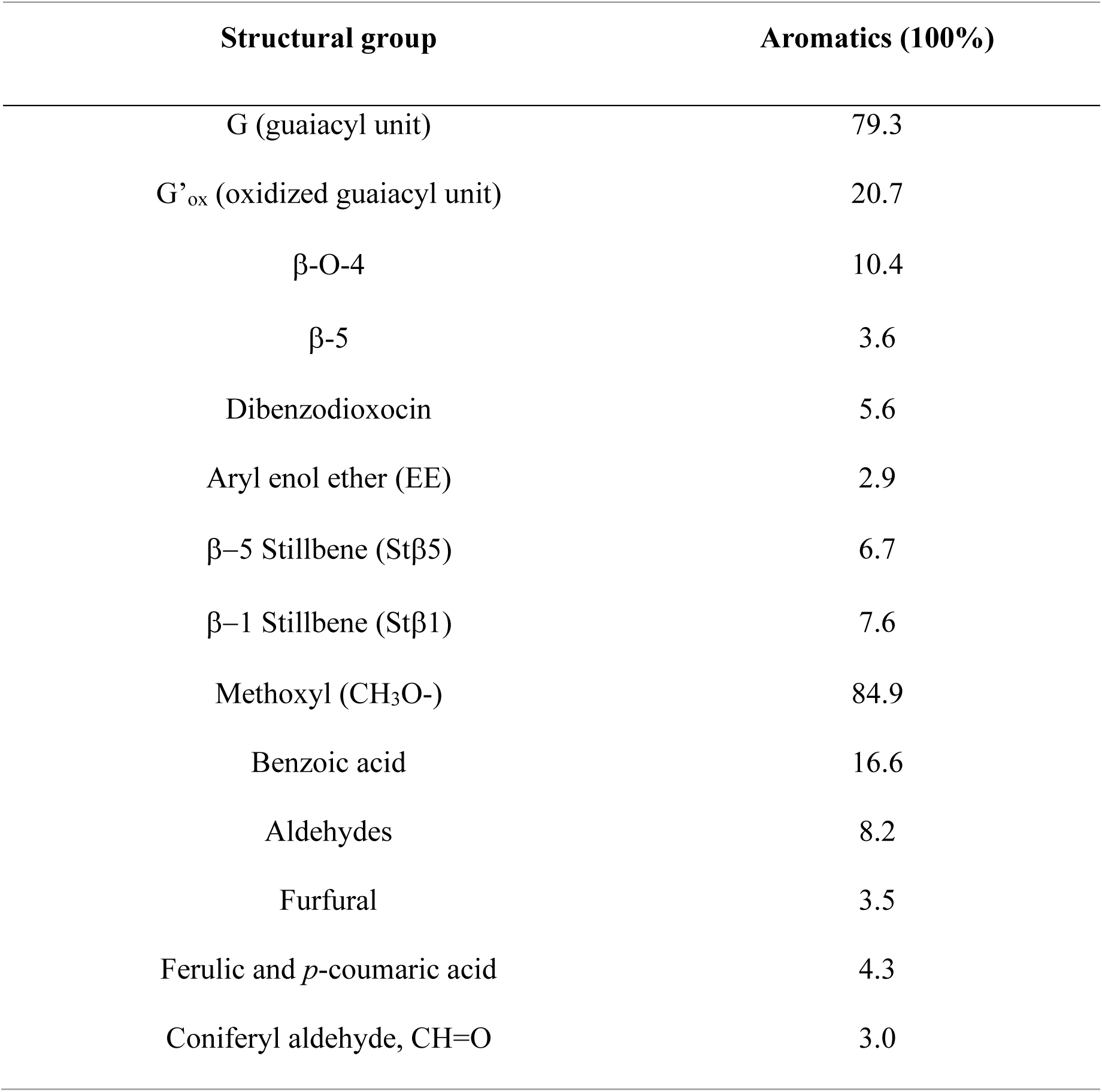
Aromatic compounds composition in the ^1^H-^13^C HSQC spectrum of the LF.

| Structural group | Aromatics (100%) |
| --- | --- |
| G (guaiacyl unit) | 79.3 |
| G'ox (oxidized guaiacyl unit) | 20.7 |
| β-O-4 | 10.4 |
| β-5 | 3.6 |
| Dibenzodioxocin | 5.6 |
| Aryl enol ether (EE) | 2.9 |
| β-5 Stillbene (Stβ5) | 6.7 |
| β-1 Stillbene (Stβ1) | 7.6 |
| Methoxyl (CH<sub>3</sub>O-) | 84.9 |
| Benzoic acid | 16.6 |
| Aldehydes | 8.2 |
| Furfural | 3.5 |
| Ferulic and *p*-coumaric acid | 4.3 |
| Coniferyl aldehyde, CH=O | 3.0 |

Semi-quantitative ^1^H-^13^C HSQC NMR analysis further revealed that the oligomeric LDMOs within the LF retained key interunit linkages, such as β-O-4 (10.4%) and β-5 (3.6%) structures. Compared to native spruce lignin, which contains over ∼50% β-O-4 linkages, these values indicate substantial modifications to the SSH lignin structure by the PHWE process as aryl enol ether (2.9%) and significan amount of benzoic acid (16.6%) has been found in LF (Table 4).^48,49^ Also, these linkages are generally recalcitrant to microbial degradation and often require microbial cell factories such as *Sphingobium* sp. SYK-6, *Dichomitus squalens,* with specialized metabolic pathways, including β-aryl ether cleavage and phenylcoumaran catabolism.^50–53^ Notably, the dibenzodioxocin linkages (5.6%), which contain stable 5-5’ carbon-carbon bonds forming lignin branching units^54^, were also identified in the LF. The side-chain region of the ^1^H-^13^C HSQC spectrum further indicated the presence of ethylenic linkages (C=C bond-containing aromatic structures), including various stilbenes (14.3%; Stβ1, Stβ5), terminating coniferyl aldehyde groups (3.0%) and ferulic or *p*-coumaric acid groups (4.3%). The stability of dibenzodioxocin and ethylenic linkages significantly influences downstream microbial depolymerization pathways, as 5-5’ dibenzodioxocin linkages and ethylenic linkages represent some of the most recalcitrant C-C bonds^52,55–57^. Notably, cleavage of these robust linkages has been observed in the anaerobic fungus *Neocallimastix californiae*.^55^ Furthermore, *Sphingomonas paucimobilis* TMY1009 and *Sphingobium* sp. SYK-6 bacterial species have demonstrated the capacity to catalyze the oxidative cleavage of ethylenic stilbene linkages.^56,58^ Additionally, microbes such as *Enterobacter* sp. employ ferulic acid decarboxylase to decarboxylate the side chains of coniferyl aldehyde, ferulic acid, and p-coumaric acid.^59^ Collectively, semi-quantitative lignin structural insights on linkage diversity will aid in identifying or developing tailored microbial and enzymatic strategies for targeted valorization of lignin compounds present within the hydrolysate.

In addition to structural characterization, ^1^H-^13^C HSQC NMR provided insights into the degree of methoxylation of the LF, which was found to be slightly lower than reported in literature. In this study, the methoxylation level was measured at 84.9%, whereas softwood lignin typically exhibits a methoxyl content of 90-95%.^60^ This deviation could be attributed to the presence of various non-methoxylated, *p*-OH phenyl and catechol-type extractives, particularly benzoic and *p*-OH benzoic acid, which were observed in high concentrations when analyzed using complementary GC-MS **(Figure S2 and Table S2)**. The availability of methoxylated LDMOs carries significant implications for microbial valorization, as microbes such as *Acetobacter woodi* and *Rhodococcus jostii* equipped with O-demethylase enzymes can utilize the methyl groups for biotransformation into value-added products.^11,61,62^ Overall, these ^1^H-^13^C HSQC-derived insights underscore the importance of detailed lignin profiling for guiding biomass selection and optimizing microbial valorization strategies.

To complement the structural and methoxylation data from ^1^H-^13^C HSQC NMR, ^31^P NMR spectroscopy was employed to selectively quantify hydroxyl group types, expanding chemical characterization of the LF. A total hydroxyl content of 7.1±0.4 mmol/g was determined in the LF. This was comprised of phenolic hydroxyl (3.6±0.4 mmol/g), aliphatic hydroxyl (1.8±0.1 mmol/g), and carboxylic hydroxyl (1.7±0.1 mmol/g) **(Figure 2)**. A range of phenolic hydroxyl groups, including condensed and uncondensed guaiacyl, *p*-hydroxyphenyl hydroxyl, and catecholic hydroxyl groups, was detected **(Figure 2)**. ^31^P NMR provides functional group-level resolution beyond the conventional methods such as HPLC and GC-MS, particularly in quantifying hydroxyl functionalities in LDMO mixtures.^63,64^ This resolution is especially valuable when assessing the suitability of biomass hydrolysates for bio-based applications, as both the type and abundance of hydroxyl groups influence lignin’s reactivity, solubility, and conversion potential.^63,65.^ Incorporating ^31^P NMR into bioprocess monitoring further enables the detection of chemical transformations within lignin, such as methoxyl-to-hydroxyl shifts, oxidative modifications, and condensation reactions, thereby improving mass balance assessments and yield predictions in lignin valorization workflows. ^1^H-^13^C HSQC, along with ^31^P NMR, delivers a robust, multidimensional view of LDMO structure and functionality, with direct relevance to targeted valorization of lignin-rich hydrolysate fractions.

**Figure 2.**
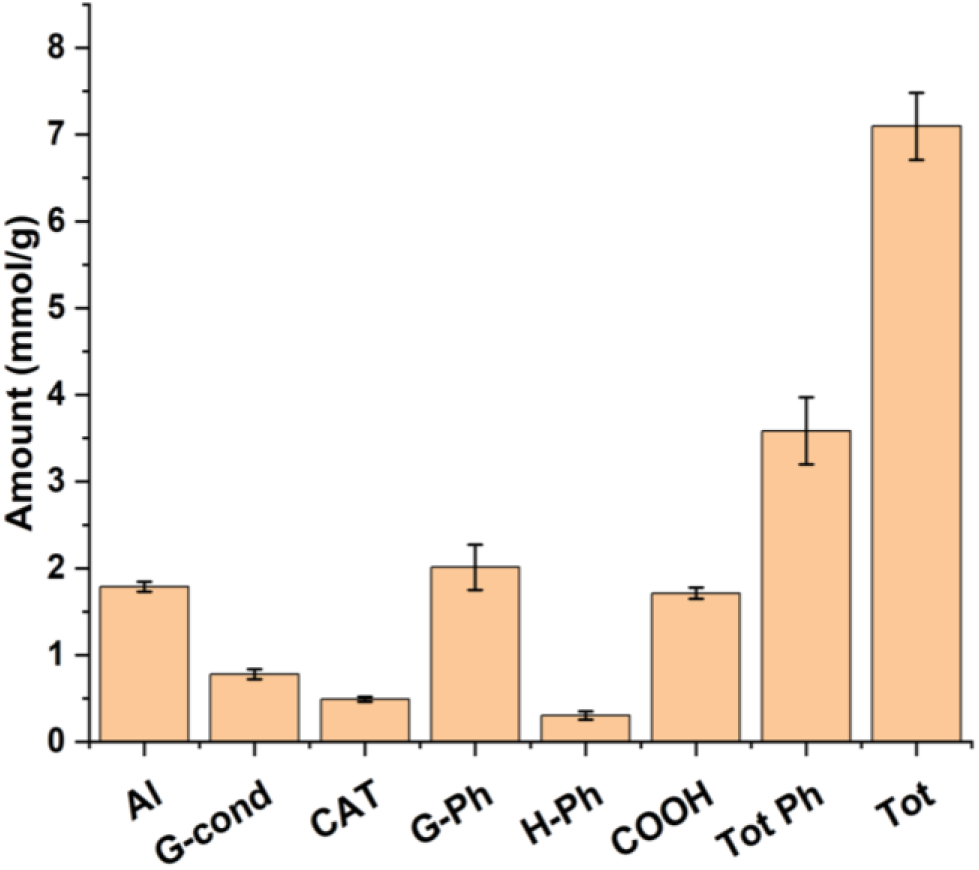
Hydroxyl group distribution in the LF quantified by ^31^P NMR (mmol/g). The quantified groups include Al (aliphatic hydroxyl), G-cond (condensed guaiacyl-type phenolics), CAT (catecholic hydroxyl), G-Ph (uncondensed guaiacyl hydroxyl), H-Ph (*p*-hydroxyphenyl hydroxyl), COOH (carboxylic acid groups), Tot Ph (sum of all phenolic hydroxyl groups), and Tot (total hydroxyl group content). The figure presents the averaged values and standard deviations (error bars) from duplicate samples.

In addition to NMR-based structural and functional group analysis, FTIR spectroscopy was employed to gain complementary molecular-level insights into the LF. The FTIR spectrum **(Figure 3)** qualitatively identified characteristic functional groups and bonding environments within LDMOs, supporting the findings from ^1^H-^13^C HSQC and ^31^P NMR. A prominent absorption band at 1705 cm^-1^, corresponding to C=O stretching vibrations from carbonyl-containing functional groups such as aldehydes, carboxylic acids, and ketones, typical oxidation products of depolymerized lignin, was identified.^66^ These observations align with the prominent aldehyde and carboxylic acid signals identified through NMR and GC-MS analyses **(Figures 1-2, Figure S2, and Table S2)**. Characteristic aromatic skeletal vibrations, including C=C stretching (observed at 1514 cm^-1^ and 1602 cm^-1^) and aromatic C-H deformation bands (observed at 1425 cm^-1^), indicated the presence of lignin aromatic backbones.^67,68^ Aliphatic and aromatic C-H stretching bands show as a wide signal at 2900-3000 cm^-1^ and additionally at 1451 cm^-1,^ indicating aliphatic C-H bending, confirming the presence of aliphatic side chains.^66^ Notably, bands at 1240 cm^-1^ and 1270 cm^-1^, assigned to aromatic C-O stretching vibrations in guaiacyl rings, confirmed the prevalence of G-type units in the LF^69^, which was in agreement with the ^1^H-^13^C HSQC findings **(Table 4).** The 1210 cm^-1^ absorption band, attributed to aryl-ether and ester linkages (C-C, C-O, C=O stretching), also indicates the β-O-4 products typically found in the LF of SSH.^70^ The hydroxyl O-H stretching vibrations of aliphatic, phenolic, and carboxylic groups show at 2400-3600 cm^-1^ as a wide band due to strong hydrogen bonding. Further, the presence of the band at 1125 cm^-1^, indicating C-O stretching in aliphatic alcohols, suggests hydroxylated side chains or residual methoxyl groups.^71^ The 1030 cm^-1^ band corresponds to C-O vibrations from both carbohydrate residues and lignin-derived alcohols, reflecting partial hemicellulose degradation and lignin-carbohydrate complex remnants.^68^ Bands between 625-900 cm^-1^ (817-832, 837, 853-858, 912 cm^-1^), indicative of C-H out-of-plane bending, further confirm the various aromatic substitution patterns of the sample.^20,72^

**Figure 3.**
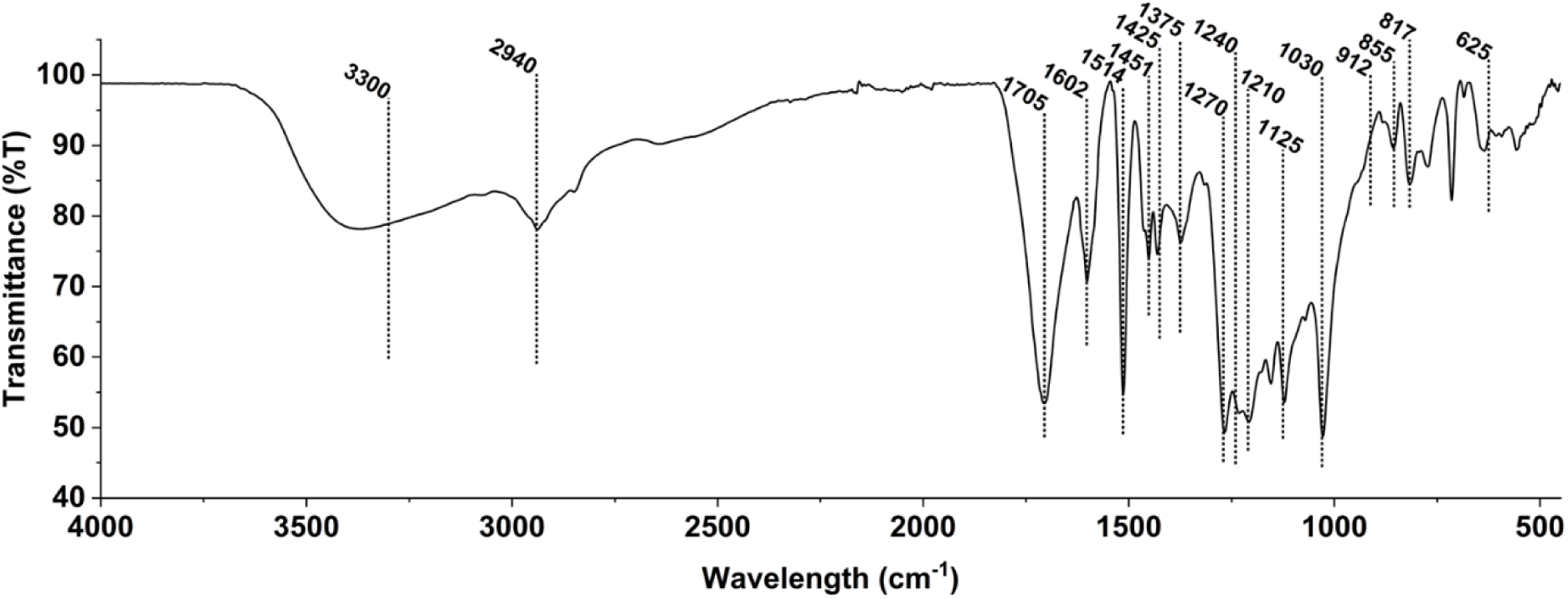
FTIR spectrum of the LF with key absorption bands corresponding to various functional groups.

FTIR analysis confirmed the presence of guaiacyl-type aromatics, oxidized LDMOs, diverse alcohols, acids, ester and aryl-ether linkages, supporting the observed structural data from NMR analyses **(Figures 1-3)**. The wide and intense carbonyl absorption band further underscores the prevalence of various oxidized products in the LF. These observations highlight the utility of FTIR as a rapid and non-invasive tool that could be used in online or at-line applications for hydrolysate uptake monitoring in downstream valorization bioprocesses.

### Quantitative Profiling of LDMOs Using Chromatographic Techniques

While NMR and FTIR spectroscopy revealed structural motifs, functional groups, and overall chemical complexity of the LF, the functional group-specific nature of such techniques limits comprehensive characterization of the diverse individual compounds in lignin-derived molecules and extractives.^73^ To overcome this limitation and achieve a more comprehensive molecular-level understanding of lignin-derived molecules and extractives, GC-MS analysis with N, O-Bis(trimethylsilyl)trifluoroacetamide (BSTFA) derivatization of LF was used to enhance volatility and facilitate the detection of lignin-derived compounds and extractives.^74^

GC-MS analysis of silylated LF detected 170 peaks, of which 83 (including two internal standards and three peaks) were identified as aromatic compounds, while the remaining 87 corresponded to fatty acids, terpenoids, alcohols, and other co-extracted extractives **(Figure S2 and Table S3)**. GC-MS data showed well-resolved chromatograms **(Figure S2)**, eliminating the need for peak deconvolution typically required for complex mixtures. Of the 80 aromatic peaks detected in the LF, 38 were identified and quantified as lignin derived monomers and extractives, representing 75% of the total chromatographic area **(Table S3)**. Major compounds detected were benzoic acid (2.7±0.2 g/L), vanillin (447.9±57.4 mg/L), 3-vanillylpropanol (414.9±16.8 mg/L), 4-hydroxy-3-methoxyphenylglycol (297.7±117.1 mg/L), coniferyl aldehyde (246.1±49.0 mg/L), and vanillic acid (234.4±15.8 mg/L). The detected high concentrations of aldehydes and carboxylic acids were consistent with the ^1^H-^13^C HSQC and ^31^P NMR data **(Figure 2** **and** **Table 3)**, corroborating the presence of oxidized aromatic structures in the LF. The elevated concentrations of these compounds likely arise from the lignin side-chain oxidation and degradation, and the degradation of hemicellulosic components during the PHWE process.^75,76^ The quantified lignin derived monomers and extractives by GC-MS accounted for 12.5% (w/w) of the total aromatics, with molar masses ranging from 110.1 to 226.2 g/mol. This observation was further supported by the molecular weight distribution profile obtained from GPC analysis of the LF **(Figure 4).**

**Figure 4.**
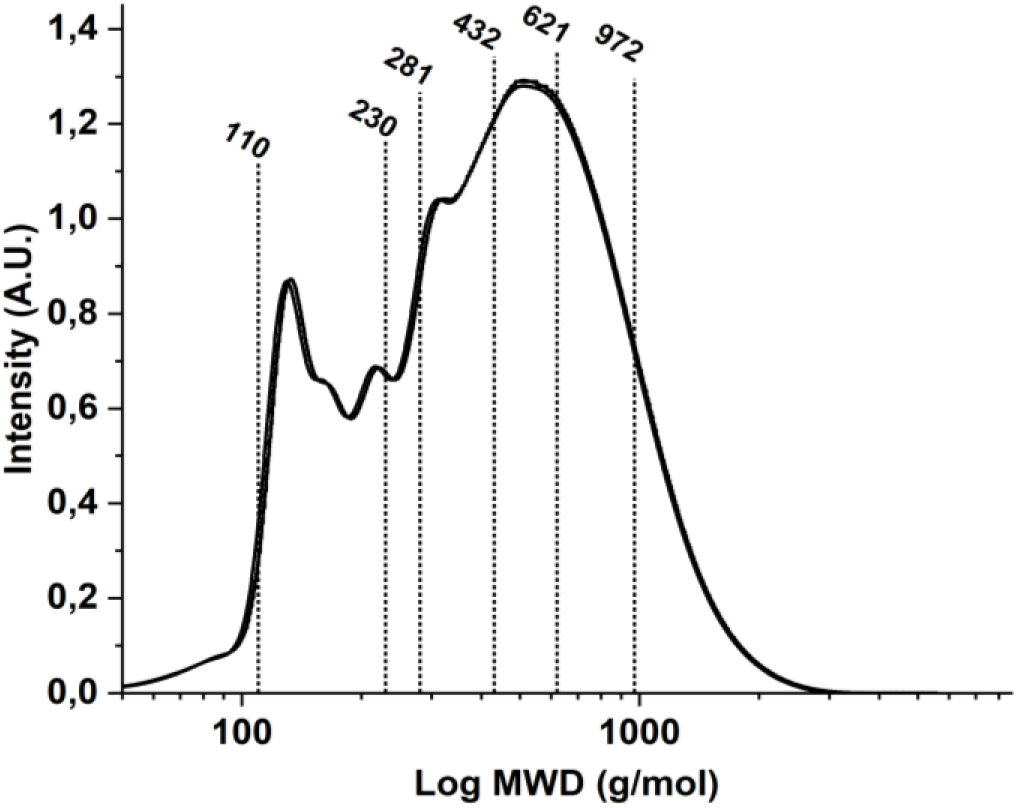
Molecular weight distribution profile of LDMOs analysed by GPC. The most significant molecular weight percentage distributions are shown.

Despite the successful quantification of 38 lignin derived monomers and extractives, GC-MS peak analysis of such complex mixtures remains challenging. Accurate identification in silylated samples is constrained by incomplete spectral libraries, overlapping retention times of structurally similar compounds, and the low confidence associated with spectral similarity.

While two-dimensional gas chromatography coupled with time-of-flight mass spectrometry (GC×GC-TOF-MS) offers improved resolution, it generates extensive datasets requiring advanced deconvolution algorithms and expert interpretation, making it cumbersome for routine quantitative analyses.^19,77^ On the other hand, pyrolysis GC-MS offers only semi-quantitative data and has lower selectivity compared to GC-MS.^3^ In contrast, the GC-MS approach enables targeted and efficient monitoring of relevant peaks (e.g., key methoxylated aromatics) pertinent to microbial bioconversion.^78^. This technique can also detect and quantify compounds such as fatty acids, terpenes, and other extractives^79.^ Although majority of extractives were not quantified here, their chromatographic area percentages were reported to provide context for their relative abundance **(Table S3).** Given the excellent linearity for lignin derived monomers and extractives detection by GC-MS (R^2^ > 0.99; **Table S2)**, an additional flame ionization detector (FID) was not necessary. Although FID is traditionally preferred for quantification due to its broader dynamic range^80^, our results demonstrate that GC-MS provides robust quantitative performance, facilitating simultaneous qualitative and quantitative analysis in a single platform. Nonetheless, the clarity of separation and reproducibility observed **(Figure S2 and Table S2-S3)** highlight the potential of GC-MS as a powerful tool for quantitative and structural characterization of lignin-derived monomers and extractives, as well as for real-time profiling of microbial utilization of hydrolysate-derived aromatics.

While GC-MS with BSTFA derivatization of LF samples enables identification and quantification of lignin-derived monomers and extractives, it cannot comprehensively detect higher molecular weight oligomers, which are typically non-volatile and poorly suited to derivatization under standard GC-MS conditions. Although comprehensive two-dimensional liquid chromatography (LC×LC) coupled with high-resolution multi-stage tandem mass spectrometry (HRMS^n^) enables analysis beyond volatility limits, it has mainly been applied to monomer identification, and quantification of oligomers remains limited due to the lack of authentic standards.^19,77^ To address this, GPC was used to estimate the molecular weight distribution of lignin oligomers. GPC revealed that 13% of the LF (w/w) comprised compounds within the 110-230 g/mol range **(Figure 4)**, which aligns with the 12.5% of lignin derived monomers and extractives detected by GC-MS **(Table S2)** in the same range, thus demonstrating consistency between the two approaches.

The molecular weight distribution indicated that 30%, 50%, 70%, and 90% of the LF (w/w) had molecular weights below 280 g/mol, 432 g/mol (median), 621 g/mol, and 972 g/mol, respectively. These values suggest the presence of lignin structures ranging from dimers to pentamers, with compounds having a molecular weight of ≥972 g/mol likely corresponding to higher-order oligomeric structures.^81–83^ The number average molecular weight, weight average (Mw), and z-average of the tested LF were 309 g/mol, 501 g/mol, and 757 g/mol, respectively, with a moderate dispersity of 1.6±0.0 This distribution indicates a dominance of low molecular weight oligomers that are more bioavailable and readily metabolized by microorganisms. For example, *Pseudomonas putida* KT2440 has been shown to catabolize oligomers with Mw values of 150, 500, and 1300 g/mol, with an optimum ∼500 g/mol.^84^ Corroborating literature has also reported that low molecular weight LDMOs are particularly suitable for the biological conversion pathway.^11,61,85^ These findings underscore the potential of the analyzed biomass-based hydrolysate for bioconversion and lignin valorization.

To complement the structural and quantitative analyses presented in this study, a complete compositional profile of the SSH, normalized to 100%, is available in **Table S4**. Overall, the integration of spectroscopic and chromatographic techniques enabled comprehensive profiling of sugars, microbial inhibitors, lignin, and extractives, yielding robust evidence for the hydrolysate’s bioconversion potential. Importantly, this multi-analytical framework is flexible and can be tailored to meet the various study requirements depending on the depth and scope of characterizations.

## CONCLUSION

This study presents an integrated multi-analytical framework for comprehensive characterization of lignocellulosic hydrolysates. By combining advanced spectroscopic and chromatographic methods, we achieved detailed profiling of elemental composition, sugars, organic acids, aromatics, lignin structural and functional groups and extractives in PHWE spruce sawdust hydrolysate. The developed workflow delivers critical insights beyond single-component analysis offers broad applicability for lignocellulosic hydrolysate profiling, establishing a foundation for future biomass valorization studies.

## SUPPORTING INFORMATION

Supporting information: The file includes additional experimental details, tables and figures. Experimental details include text on chemicals used, sample preparation, extraction of lignin derived monomers and oligomers, and calculation of GC-MS response factors for quantitative analysis. Tables present the linearity and retention times for sugars, inhibitors, and lignin derived monomers and extractives standards, as well as peak integration and hydrolysate composition. Figures present the ^1^H NMR spectrum of extracted lignin fractions and GC-MS chromatograms of derivatized lignin fraction with internal standards.

## Supporting information

The file includes additional experimental details, tables and figures

## ACKNOWLEDGMENTS

The authors acknowledge Research Engineer Ilkka Välimaa for his contribution in ICP-OES experimentation and data analysis. This research was financially supported by the Assistant Professor startup research grant (Aalto University) and the Research Council of Finland (Grant Number 353673) awarded to R.M. The Research Council of Finland’s FinnCERES (Competence Center for Materials Bioeconomy) flagship programme (grant number 345553) is also gratefully acknowledged. The authors are grateful to Jaakko Pajunen and Boreal Bioproducts for their support and for providing the hydrolysate used in the study

