## Supplementary material for "Integrated Multi-analytical Framework for Comprehensive Characterization of Lignocellulosic Hydrolysates": The file includes additional experimental details, tables and figures

Table of Contents

Supporting materials and methods ....S2

Chemicals used for sample preparation and analytics ....S2

Extraction of lignin derived monomers and oligomers ....S3

Sample preparation ....S3

Calculation of GC-MS response factors and quantitative determination of assigned monomers ....S5

Supporting Results ....S5

Table S1: Linearity of detection and retention time of SSH sugars and inhibitors detected in HPAEC-PAD and HPLC, respectively. Acetic acid, formic acid and levulinic acid at 210 nm, while furfural and 5-hydroxymethylfurfural (HMF) at 254 nm UV detection was used for data analysis. ....S5

Figure S1. $^{1}$H NMR spectrum of the LF in pressurized-hot-water extracted SSH. ....S7

Figure S2. GC-MS chromatogram of the silylated LF. Sub-figures A- D presents the 170 peaks detected. Peaks corresponding to the internal standards 4-hydroxyacetophenone and veratric acid are labeled as 42 and 78, respectively. ....S9

Table S2: Linearity of detection of standards and concentration of lignin derived monomers and extractives detected by GC-MS analysis. The total number of calibration points for standards was 4....S9

Table S3: GC-MS Peak Integration: Peak Number, Area (%), and Similarity Index....S13

Table S4: SSH composition (g per 100 g Dry Biomass$^{\Delta}$ (DW)). ....S21

Reference ........................................................................................................................S22

### Supporting materials and methods

#### Chemicals used for sample preparation and analytics

Sulfuric acid (98%), sodium hydroxide (1M), sulfuric acid (72%), methanol (≥99.9%), nitric acid (67-69%), sodium chloride (≥99%), anhydrous magnesium sulfate (99.5%), ethyl acetate (EtOAc, ≥99.5%), L-(+)-arabinose (≥99%), L-rhamnose monohydrate (≥99%), D-galactose (≥99%), D-(+)-glucose (≥99.5%), D-(+)-xylose (≥99%), D-(+)-mannose (≥99%), acetic acid (≥99.7%), formic acid (≥98%), levulinic acid (98%), 5-hydroxymethylfurfural (HMF, ≥99%), and furfural (≥99%), dimethylformamide (DMF, anhydrous 99.8%), pyridine (99.8%), tetrahydrofuran (THF, ≥99.9%), dichloromethane, N-hydroxy-5-norbornene-2,3-dicarboxylic acidimide (n-NHDI, 97.0%), chromium (III)acetylacetonate ($Cr(acac)_3$, ≥98.0%), phosphitylating reagent 2-chloro-4,4,5,5-tetramethyl-1,3,2-dioxaphospholane (95%), N, O-bis (trimethylsilyl) trifluoroacetamide with 10% TMS-Cl (BSTFA; GC derivatization grade 98%), 4-hydroxybenzoic acid (≥99%), benzoic acid (≥99.5%), homovanillyl alcohol (≥99%), 4-hydroxyacetophenone (≥95%), 4-hydroxybenzyl alcohol (≥98%), 3,4-dihydroxybenzoic acid (≥97%), vanillin (≥99%), veratric acid (≥99%), 3,4-dihydroxybenzaldehyde (≥97%), and *p*-coumaric acid (≥98%), 3-(4-hydroxyphenyl)-1-propanol (99%), vanillic acid (≥97%), homovanillic acid (≥99%), 4-hydroxy-3-methoxymandelic acid (≥98%), eugenol (99%), syringaldehyde (≥98%), 4-hydroxy-3-methoxyacetophenone (98%), ferulic acid (≥99%), catechol (≥95%), and 2-hydroxy-3-methoxybenzoic acid (97%) were purchased from Sigma-Aldrich (Finland). Analytical grade deuterated dimethyl sulfoxide (DMSO-$d_6$) and deuterated chloroform ($CDCl_3$, 99.8%) were obtained from EURISO-TOP (France). All chemicals were used as received, except for dimethylformamide and pyridine, which were dried under molecular sieves (3Å) prior to use.

Extraction of lignin derived monomers and oligomers

To ensure optimal extraction conditions for quantitative and qualitative analyses of phenolic compounds, soluble lignin was extracted from 10 mL of spruce sawdust hydrolysate (SSH) using ethyl acetate (EtOAc) as a solvent after pH adjustment to 2 using 32% HCl. A liquid-liquid extraction was performed in a conical separating funnel by EtOAc addition in a 3:1 ratio (EtOAc:sample) and using 30 mL brine to improve extraction of water-soluble compounds into the organic phase. The mixture was shaken thoroughly to facilitate extraction, and the aqueous phase was separated and re-extracted twice yielding a total of three extractions with a total volume of 100 mL EtOAc. The combined organic phase was washed with 20-30 mL of brine to remove residual water. Anhydrous magnesium sulfate was then added to further remove water from the organic phase and the obtained solution was filtered through a funnel packed with a cotton plug and complete solvent removal was carried out using a rotary evaporator (Rotavapor Büchi R-210) at 133 mbar pressure, 6 rpm in a water bath temperature of 40°C. The organic phase, referred to as lignin fraction (LF) (43.3±1.2 g/L, 74 % w/w of total aromatics in SSH), was dried in vacuum-oven to eliminate residual EtOAc before further analysis. Samples were stored in a desiccator before analysis.

Sample preparation

To determine the total sugar concentration in SSH, the two-step acid hydrolysis procedure described in the NREL Laboratory Analytical Procedure (LAP), document NREL/TP-510-42618, was employed. Following hydrolysis, the monosaccharide concentrations were quantified using High-Performance Anion-Exchange Chromatography (HPAEC) coupled with Pulsed Amperometric Detection (PAD). The oligomeric sugar content was then calculated by subtracting the native monomeric sugar concentrations present in the untreated SSH.

For the determination of inorganic salts in the SSH, 300 µL of the SSH was transferred into a 50 mL Teflon microwave digestion vessel. Then, 10 mL of nitric acid was slowly added under a fume hood. The vessel was sealed with a lid and placed in a microwave oven, where the contents were heated to 180°C and maintained at that temperature for 15 minutes. After cooling, the vessel was cautiously opened under the fume hood. The digested sample was then transferred to a 50 mL volumetric flask and diluted to volume with deionized water. Finally, the samples were filtered and further diluted as necessary to achieve the appropriate concentrations for inductively Coupled Plasma - Optical Emission Spectroscopy (ICP-OES) analysis.

For $^{1}H$ and HSQC NMR (Nuclear Magnetic Resonance), 70 mg vacuum oven-dried samples were dissolved in 700 µL of deuterated dimethyl sulphoxide (DMSO-$d_6$) before analysis.

Sample preparation for $^{31}P$-NMR analysis was done following the method described by Maltari et al.[1] Vacuum oven-dried lignin (30 mg) was accurately weighed into a 2 mL glass vial containing a mini magnetic stirrer. Phosphitylation was performed by sequentially adding anhydrous solvents and solutions: 150 µL DMF, 100 µL pyridine, 200 µL internal standard (9.27 mg/mL nNHDI in $CDCl_3$), 50 µL $Cr(acac)_3$ relaxation agent (5.6 mg/mL in $CDCl_3$), 300 µL $CDCl_3$, and 150 µL phosphitylating reagent 2-chloro-4,4,5,5-tetramethyl-1,3,2-dioxaphospholane under nitrogen flow. The sample was dissolved by stirring for 10 minutes, then transferred to an NMR tube, and the $^{31}P$ NMR spectrum was recorded within 1 hour of sample preparation.

To quantify lignin derived monomers by using Gas Chromatography-Mass Spectrometry (GC-MS), the vacuum oven dried samples were derivatized via BSTFA to enhance volatility and stability of lignin derived monomers and oligomers. Each sample (3 mg) for analysis was prepared by adding 100 µL of dimethyl formamide (DMF), 10 µL of pyridine, and 50 µL of

10% BSTFA. The mixture was heated at 65°C for 15 minutes, with occasional shaking to ensure complete derivatization. Following the reaction, the samples were filtered through a 0.22 µm syringe filter and transferred into GC vials for analysis.

For Gel Permeation Chromatography (GPC) analysis a calibration curve was established using toluene and polystyrene standards of known molecular weights. Both the samples and calibration standards were prepared in THF at a concentration of 2 mg/mL, followed by filtration through 0.22 µm syringe filters to remove particulates prior to injection.

Calculation of GC-MS response factors and quantitative determination of assigned monomers

The response factor (*F*) of each component in calibration mixture respect to sample were calculated using following equation:

$$F = \frac{A_{is}}{A_i} \quad (1)$$

Where $A_{is}$ is chromatogram peak area of internal standard in calibration mixture, $A_i$ is peak area of internal standard in sample for same concentration.

Concentration of each compound ($C_F$) in the sample using response factor calculated using following equation:

$$C_F = C_S \times F \quad (2)$$

Where $C_s$ is concentration of each compound calculated using corresponding standard in the sample; *F* is the integration factor applied to neutralize differences in the internal standard signal between the standard mixture and the sample.

### Supporting Results

**Table S1:** Linearity of detection and retention time of SSH sugars and inhibitors detected in HPAEC-PAD and HPLC, respectively. Acetic acid, formic acid and levulinic acid at 210 nm, while furfural and 5-hydroxymethylfurfural (HMF) at 254 nm UV detection was used for data analysis.

| | | Number of Calibration Points | Linearity of detection ($R^2$) | Retention time (min) |
|---|---|---|---|---|
| Sugars | Arabinose | 4 | 0.998 | 13.79 |
| | Rhamnose | | 0.998 | 15.13 |
| | Galactose | | 1 | 18.43 |
| | Glucose | | 1 | 23.19 |
| | Xylose | | 0.999 | 28.02 |
| | Mannose | | 1 | 31.3 |
| Inhibitors | Acetic Acid | 4 | 1 | 17.1 |
| | Formic Acid | | 1 | 18.6 |
| | Levulinic acid | | 1 | 19.98 |
| | HMF | | 0.997 | 37.29 |
| | Furfurals | | 1 | 53.47 |

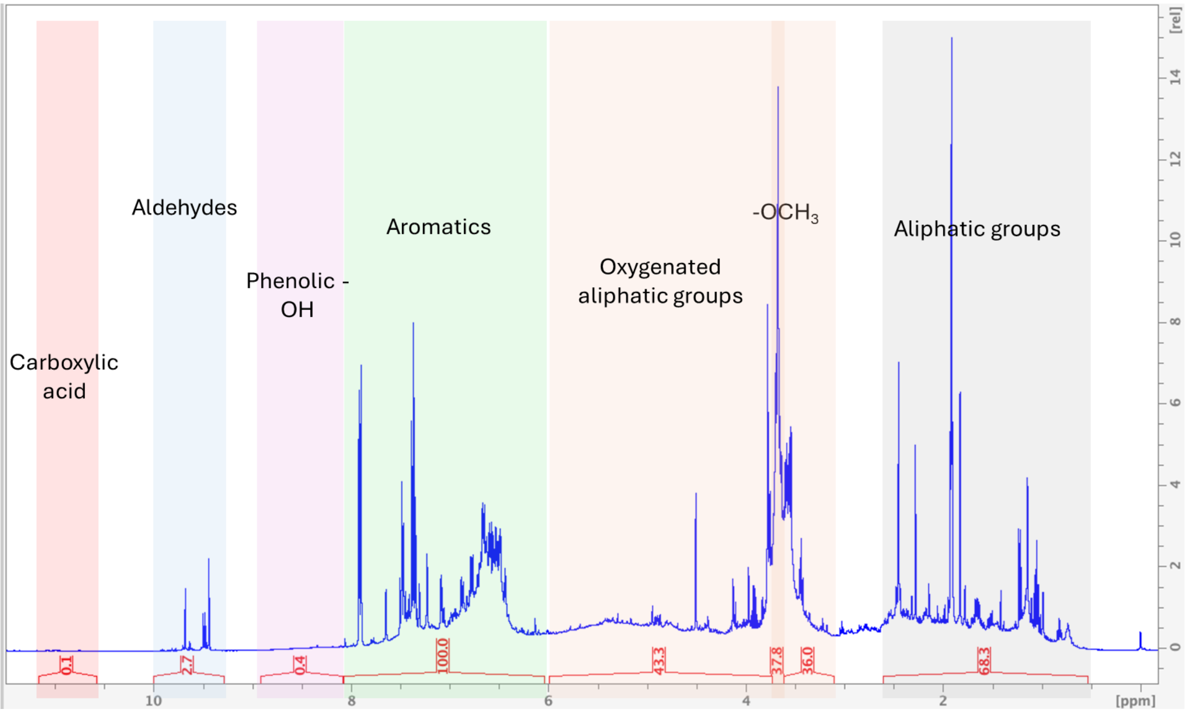

**Figure S1.** [1]H NMR spectrum of the LF in pressurized-hot-water extracted SSH.

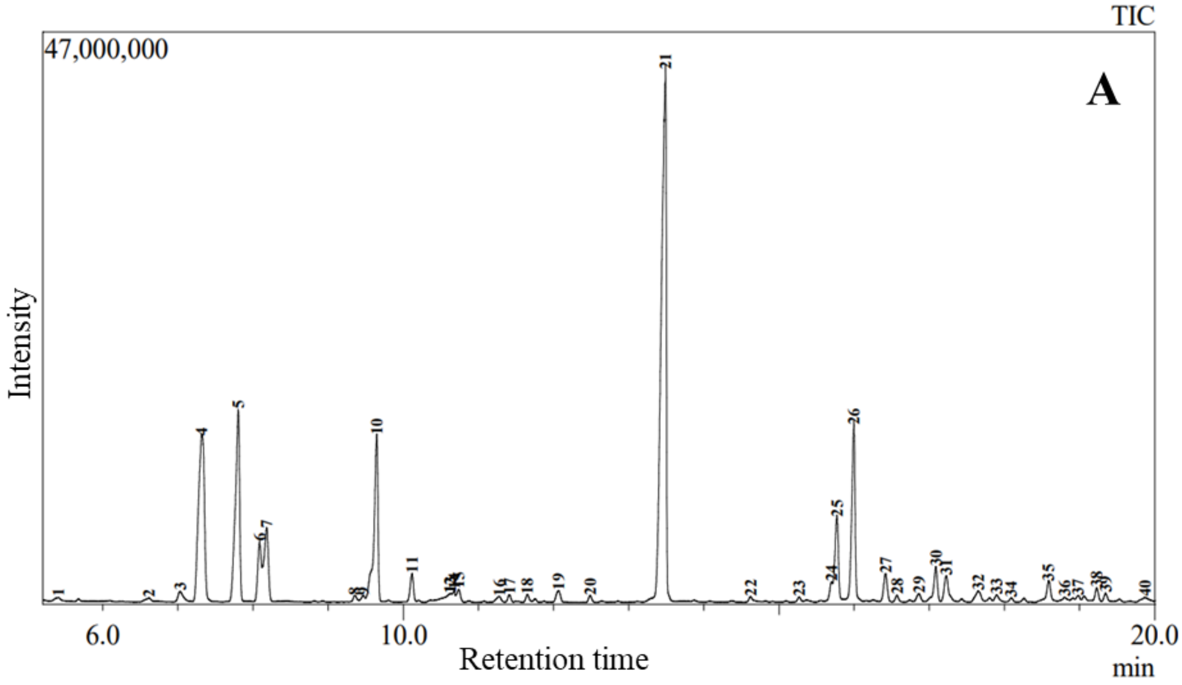

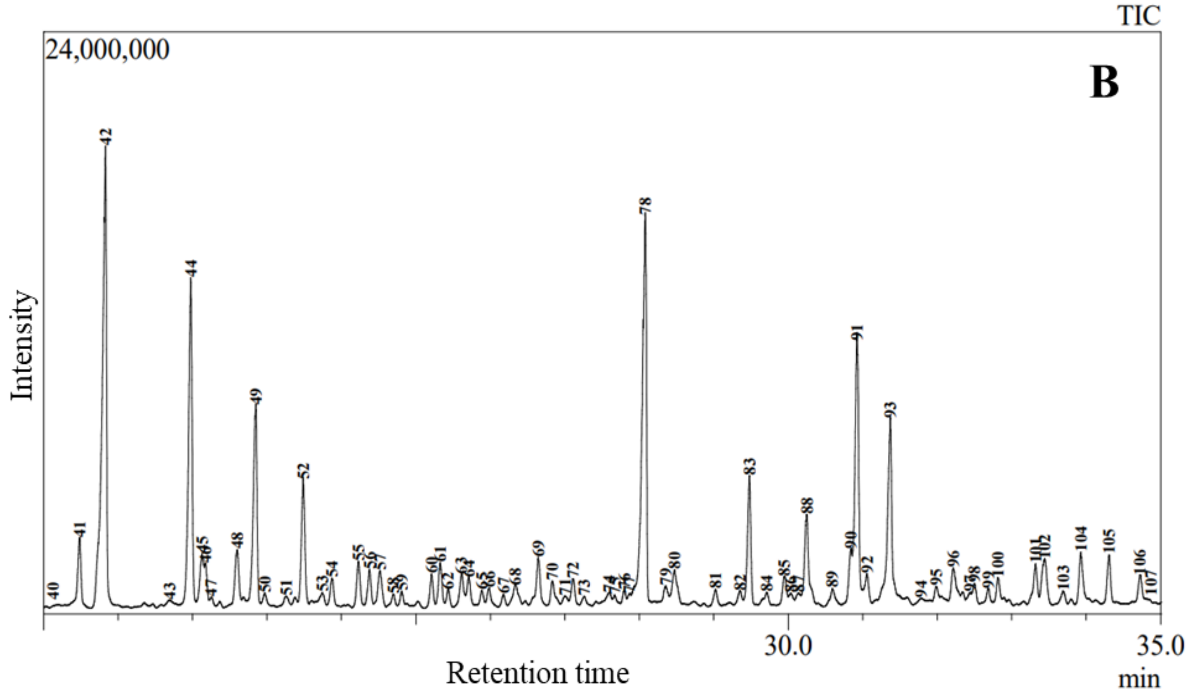
TIC
24,000,000
B
Intensity
30.0
35.0
Retention time
min

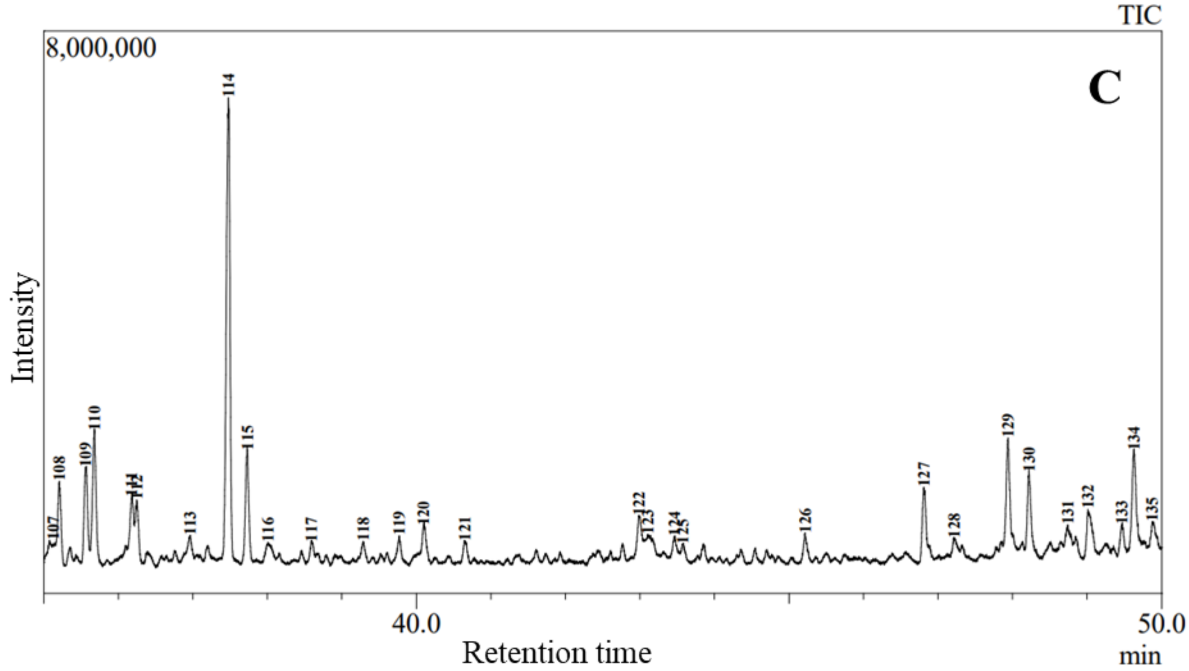
TIC
8,000,000
C
Intensity
40.0
50.0
Retention time
min

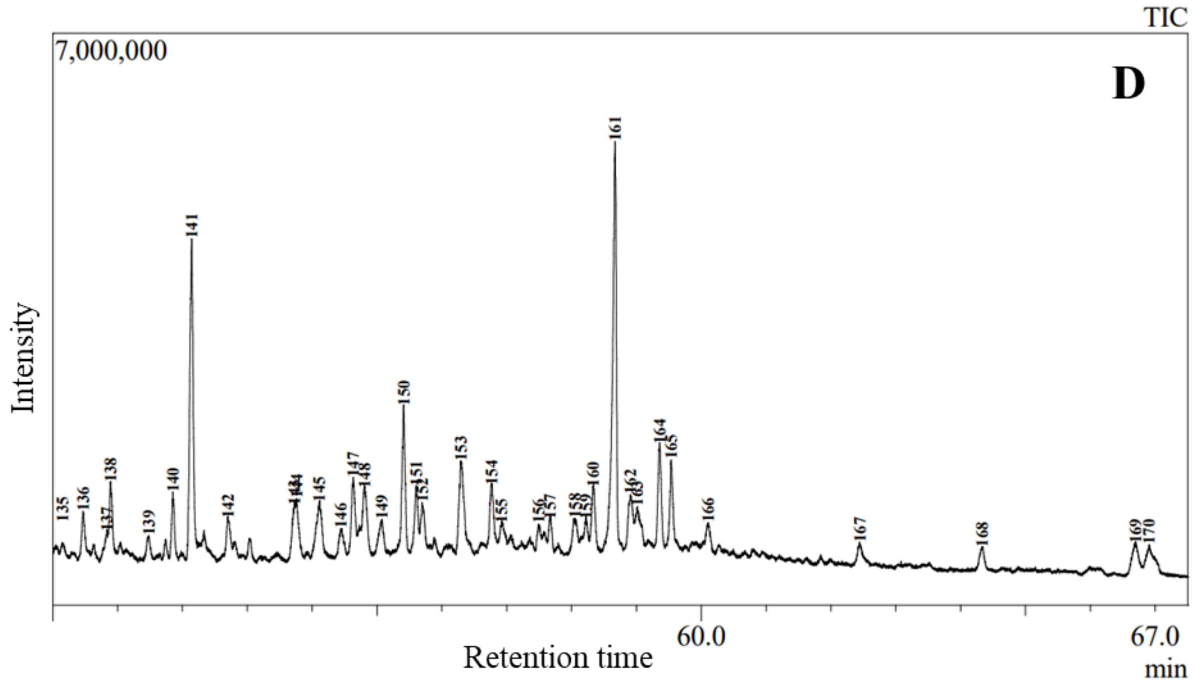

**Figure S2.** GC-MS chromatogram of the silylated LF. Sub-figures A- D presents the 170 peaks detected. Peaks corresponding to the internal standards 4-hydroxyacetophenone and veratric acid are labeled as 42 and 78, respectively.

**Table S2:** Linearity of detection of standards and concentration of lignin derived monomers and extractives detected by GC-MS analysis. The total number of calibration points for standards was 4.

| **Standards*** | **Linearity of detection ($R^2$)** | **Phenolics** | **Concentration (mg/L)** | **Peak number**[Γ] |
|---|---|---|---|---|
| 4-Hydroxybenzoic acid | 1 | 1,2-Ethanediol monobenzoate | 36.9±14.3 | 14 |
| Benzoic Acid | 1 | Benzoic Acid | 2735.7±244.0 | 21 |
| Homovanillyl alcohol | 0.999 | 1,3,5-Benzetriol | 215.0±48.9 | 25 |

| | | | | |
|---|---|---|---|---|
| 4-Hydroxyacetophenone | 0.996 | 4-Hydroxybenzaldehyde | 118.4±14.3 | 32 |
| 4-Hydroxybenzylic alcohol | 0.999 | Hydroquinone | 9.0±0.3 | 36 |
| Vanillin | 0.996 | Vanillin | 447.9±57.4 | 49 |
| 4-Hydroxybenzoic acid | 1 | 3-Hydroxybenzoic acid | 45.6±1.7 | 54 |
| 4-Hydroxyacetophenone | 0.996 | 4-Hydroxyphenylvinyl alcohol | 21.8±1.9 | 58 |
| 3,4-Dihydroxybenzaldehyde | 0.996 | 3,4-Dihydroxybenzaldehyde | 71.2±15.4 | 61 |
| 4-Hydroxybenzoic acid | 1 | 4-Hydroxybenzoic acid | 67.6±10.7 | 63 |
| 4-Hydroxybenzoic acid | 1 | 3-Hydroxyphenylacetic acid | 123.1±28.5 | 69 |
| *p*-Coumaric acid | 1 | p-Hydroxyphenylacrylic acid | 90.9±32.1 | 72 |

| | | | | |
|---|---|---|---|---|
| *p*-Coumaric acid | 1 | o-Coumaric acid | 59.4±27.3 | 79 |
| 3-(4-Hydroxyphenyl)-1-propanol | 0.999 | 2-Hydroxyphenethyl alcohol | 18.7±0.8 | 82 |
| Vanillic Acid | 1 | Vanillic Acid | 234.4±15.8 | 83 |
| Homovanillic Acid | 0.996 | Homovanillic Acid | 29.6±3.3 | 84 |
| 4-Hydroxy-3-methoxymandelic acid | 0.996 | 3-Methoxylsalicylic acid | 128.9±52.6 | 88 |
| Homovanillyl alcohol | 0.999 | 3-Vanilpropanol | 414.9±16.8 | 91 |
| 3,4-Dihydroxybenzoic acid | 1 | 3,4-Dihydroxybenzoic acid | 35.0±4.2 | 92 |
| Eugenol | 0.996 | Coniferyl aldehyde | 246.1±49.0 | 93 |
| 3,4-Dihydroxybenzoic acid | 1 | 3,4-Dihydroxycinnamyl alcohol | 46.6±1.5 | 96 |
| Syringaldehyde | 0.996 | 3-Hydroxy-4-methoxybenzaldehyde | 15.1±16.9 | 97 |
| 4-Hydroxy-3-methoxymandelic acid | 0.996 | 3,4-Dihydroxycinnamic acid | 17.3±1.02 | 99 |

| | | | | |
|---|---|---|---|---|
| 4-Hydroxy-3-methoxymandelic acid | 0.996 | 4-Hydroxy-3-methoxymandelic acid | 27.4±9.8 | 100 |
| 4-Hydroxybenzoic acid | 1 | 3-Methylsalicylic acid | 73.7±25.3 | 101 |
| Homovanillyl alcohol | 0.999 | trans-Coniferyl alcohol | 73.3±21.8 | 104 |
| 4-Hydroxy-3-methoxyacetophenone | 0.996 | 2-Hydroxy-4-methoxyacetophenone | 161.4±43.6 | 106 |
| Vanillic acid | 1 | 3-(4-Hydroxy-3-methoxyphenyl)propionic acid | 84.5±22.6 | 110 |
| Homovanillic Acid | 0.996 | 4-Hydroxy-3-methoxyphenylglycol | 69.9±16.8 | 111 |
| 3,4-Dihydroxybenzoic acid | 1 | 2,5-Dihydroxybenzoic acid | 7.5±6.6 | 116 |
| Ferulic acid | 0.996 | 3-(4-Hydroxy-3-methoxyphenyl)-1,2-propanediol | 26.5±3.8 | 119 |
| Homovanillyl alcohol | 0.999 | 1,2-Dihydroxy-3-(3,4-dimethoxyphenyl)propane | 23.3±3.3 | 120 |

| | | | | |
|---|---|---|---|---|
| Catechol | 0.996 | Catechol | 29.4±1.0 | 127 |
| 4-Hydroxy-3-methoxymandelic acid | 0.996 | 3-Hydroxy-3-(4'-hydroxy-3'-methoxyphenyl) propionic acid | 70.8±19.6 | 129 |
| 2-Hydroxy-3-methoxybenzoic acid | 1 | 3-Hydroxy-4-methoxy phenyl ethylene glycol | 297.7±117.1 | 130 |
| 4-Hydroxy-3-methoxymandelic acid | 0.996 | 3-Ethoxy-4-hydroxymandelic acid | 32.8±3.4 | 132 |
| 4-Hydroxy-3-methoxymandelic acid | 0.996 | Dihydroferulic acid | 76.1±18.8 | 150 |
| 4-Hydroxy-3-methoxymandelic acid | 0.996 | o-Methoxymandelic acid | 31.3±1.5 | 151 |

Γ - The peak number assigned to GC-MS chromatogram peaks in figure S1; * - Calibration standards for quantification were chosen based on either exact chemical identity or, where unavailable, the closest structural analogues. Selection criteria included matching key functional groups, similar bonding patterns inside chains, and comparable physicochemical properties such as boiling points. This approach aligns with established analytical practices to ensure reliable and accurate quantification when pure standards are limited or unavailable.[2,3]

**Table S3: GC-MS Peak Integration: Peak Number, Area (%), and Similarity Index**

| Name | Chromatographic area % | Peak number[Γ] |
|---|---|---|
| Propylene glycol | 0.12 | 1 |
| Chrysanthemyl alcohol | 0.11 | 2 |
| 1,3-Propanediol | 0.29 | 3 |
| Lactic acid | 5.49 | 4 |
| Glycolic acid | 4.53 | 5 |
| 2-Methylpropanoic acid | 1.18 | 6 |
| 3-Hydroxypropanoic acid | 1.8 | 7 |
| Furfuryl alcohol | 0.13 | 8 |
| Oxalic acid | 0.14 | 9 |
| 2-Furoic acid | 3.74 | 10 |
| 3-Hydroxypropanoic acid | 0.49 | 11 |
| Phenyl(3-tridecyloxiranyl)methanone | 0.31 | 12 |
| Benzophenone | 0.09 | 13 |
| 1,2-Ethanediol monobenzoate | 0.16 | 14 |
| 3-Heptanol | 0.26 | 15 |
| 2-Methyl-3-buten-2-ol | 0.16 | 16 |
| (E)-1,3-Cyclohexanediol | 0.12 | 17 |
| Glycerol | 0.15 | 18 |
| 2-Methylglycerol | 0.27 | 19 |
| 1-(2-Methoxy-1-methylethoxy)-2-propanol | 0.13 | 20 |
| Benzoic acid | 14.85 | 21 |
| Itaconic acid | 0.11 | 22 |
| 3-Methylene-1,4-dihydroxybutane | 0.1 | 23 |

| | | |
|---|---|---|
| Maleic acid | 0.33 | 24 |
| 1,3,5-Benzetriol | 1.7 | 25 |
| Succinic acid | 3.35 | 26 |
| 2-Hydroxyhexanoic acid | 0.56 | 27 |
| 2-Hydroxyhexanoic acid | 0.12 | 28 |
| Salicylic acid | 0.17 | 29 |
| Fumaric acid | 0.71 | 30 |
| 2-Ketobutyric acid | 0.63 | 31 |
| 4-Hydroxybenzaldehyde | 0.32 | 32 |
| 3-Methyl-5-keto-3-hexenoic acid | 0.13 | 33 |
| 1-Octanol | 0.07 | 34 |
| 2-Methylglycerol | 0.52 | 35 |
| Hydroquinone | 0.08 | 36 |
| Glutaric acid | 0.08 | 37 |
| 2-Methyl-1,2-dihydroxybutane | 0.23 | 38 |
| 2-Methyl-1,2-dihydroxybutane | 0.15 | 39 |
| 4'-Hydroxyacetophenone | 0.16 | 40 |
| Itaconic acid | 0.71 | 41 |
| 4'-Hydroxyacetophenone | 5.59 | 42 |
| Pyroglutamic acid | 0.11 | 43 |
| Malic acid | 3.28 | 44 |
| 2-Methyl-4-ketoglutaconic acid | 0.57 | 45 |
| Adipic acid | 0.3 | 46 |
| Salicylic acid | 0.08 | 47 |
| Terpin | 0.67 | 48 |

| | | |
|---|---|---|
| Vanillin | 2.1 | 49 |
| trans-Aconitic acid | 0.1 | 50 |
| 2,3-Dihydroxy-2-methylpropanoic acid | 0.11 | 51 |
| 3,6-Di-O-methylglucopyranose | 1.2 | 52 |
| 2,4-Di-O-methylglucopyranose | 0.15 | 53 |
| 3-Hydroxybenzoic acid | 0.26 | 54 |
| Terpin | 0.47 | 55 |
| 3-O-Methyl-β-D-glucopyranose | 0.38 | 56 |
| 2-Hydroxypentanedioic acid | 0.44 | 57 |
| 4-Hydroxyphenylvinyl alcohol | 0.12 | 58 |
| Adipic acid | 0.12 | 59 |
| Isoleucine | 0.32 | 60 |
| 3,4-Dihydroxybenzaldehyde | 0.46 | 61 |
| 2-Hydroxy-2-pentenedioic acid | 0.17 | 62 |
| 4-Hydroxybenzoic acid | 0.38 | 63 |
| Cyclopentane, 1-isopropylidene-2-ol | 0.29 | 64 |
| D-(-)-Ribofuranose | 0.14 | 65 |
| D-Arabinose | 0.19 | 66 |
| L-Pyroglutamic acid | 0.11 | 67 |
| L-(+)-Rhamnopyranose | 0.3 | 68 |
| 3-Hydroxyphenylacetic acid | 0.56 | 69 |
| D-Ribose | 0.27 | 70 |
| Methyl 3,4-O-isopropylidene-L-threonate | 0.1 | 71 |
| p-Hydroxyphenylacrylic acid | 0.26 | 72 |
| 2-Hydroxyhexanedioic acid | 0.1 | 73 |

| | | |
|---|---|---|
| L-(-)-Sorbofuranose | 0.2 | 74 |
| 1,8-cis-Undecadien-5-yne-3,7-diol | 0.08 | 75 |
| 1-Cyclopentene-3,5-dione, 4-hydroxy-1,2,4-triol | 0.1 | 76 |
| Terpin | 0.08 | 77 |
| Veratric acid | 4.83 | 78 |
| o-Coumaric acid | 0.21 | 79 |
| D-Xylopyranose | 0.47 | 80 |
| 2,3-Dihydroxybenzoic acid | 0.18 | 81 |
| 2-Hydroxyphenethyl alcohol | 0.16 | 82 |
| Vanillic acid | 1.2 | 83 |
| Homovanillic acid | 0.12 | 84 |
| D-Xylopyranose | 0.33 | 85 |
| 2-Hydroxy-N'-(2-(hydroxyimino)-1-phenylethylidene)benzohydrazide | 0.14 | 86 |
| 1-Undecanol | 0.15 | 87 |
| 3-Methyoxylsalicylic acid | 1.09 | 88 |
| 2-Hydroxyhexanoic acid | 0.24 | 89 |
| 4-Methyl-1,2-dihydroxypentane | 0.52 | 90 |
| 3-Vanilpropanol | 2.88 | 91 |
| 3,4-Dihydroxy-benzoic acid | 0.32 | 92 |
| Coniferyl aldehyde | 2.28 | 93 |
| Malonic acid | 0.08 | 94 |
| Methyl 4-hydroxy-3-methoxymandelate | 0.18 | 95 |
| 3,4-Dihydroxycinnamyl alcohol | 0.43 | 96 |
| 3-Hydroxy-4-methoxybenzaldehyde | 0.07 | 97 |

| | | |
|---|---|---|
| 1,3-Dihydroxyacetone dimer | 0.18 | 98 |
| 3,4-dihydroxycinnamic acid | 0.13 | 99 |
| 4-Hydroxy-3-methoxymandelic acid | 0.22 | 100 |
| 3-Methylsalicylic acid | 0.48 | 101 |
| 2-Methyl-2-(p-methoxy)mandelate | 0.67 | 102 |
| 4-Hydroxy-3-methoxyphenylglycol | 0.17 | 103 |
| trans-Coniferyl alcohol | 0.57 | 104 |
| 3-Phenyllactic acid | 0.45 | 105 |
| 2-Hydroxy-4-methoxyacetophenone | 0.34 | 106 |
| Salidroside | 0.09 | 107 |
| 4-Acetylphenol nonyl ether | 0.29 | 108 |
| Vanilpyruvate | 0.34 | 109 |
| 3-(4-Hydroxy-3-methoxyphenyl)propionic acid | 0.46 | 110 |
| 4-Hydroxy-3-methoxyphenylglycol | 0.25 | 111 |
| Palmitic acid | 0.22 | 112 |
| Palmitoleonitrile | 0.1 | 113 |
| Oxanilic acid | 1.64 | 114 |
| 2-Methyl-2-(p-methoxy)mandelate | 0.37 | 115 |
| 2,5-Dihydroxybenzoic acid | 0.11 | 116 |
| 1-(2'-Hydroxyphenyl)-1-pentanol | 0.08 | 117 |
| 1-(2'-Hydroxyphenyl)-1-pentanol | 0.08 | 118 |
| 3-(4-Hydroxy-3-methoxyphenyl)-1,2-propanediol | 0.11 | 119 |
| 1,2-Dihydroxy-3-(3,4-dimethoxyphenyl)propane | 0.19 | 120 |
| Myristic acid | 0.09 | 121 |
| Oleamide | 0.18 | 122 |

| | | |
|---|---|---|
| 3-Methoxy-4-hydroxybenzenepropanoic acid | 0.13 | 123 |
| (3R-E)-Matairesinol | 0.1 | 124 |
| 2,3-Dimercaptobutane-1,4-diol | 0.08 | 125 |
| 3-Vanil-1,2-dihydroxypropane | 0.11 | 126 |
| Catechol | 0.31 | 127 |
| 4-Methoxy-2,5-dimethylphenol | 0.08 | 128 |
| 3-Hydroxy-3-(4'-hydroxy-3'-methoxyphenyl) propionic acid | 0.41 | 129 |
| 3-hydroxy-4-methoxy phenyl ethylene glycol | 0.28 | 130 |
| 1-Butyn-4-ol | 0.08 | 131 |
| 3-Ethoxy-4-hydroxymandelic acid | 0.24 | 132 |
| Methyl 4-methoxy-3-hydroxyphenylacetate | 0.11 | 133 |
| 1-(2'-Hydroxyphenyl)-1-pentanol | 0.43 | 134 |
| 3-Methoxymandelic acid | 0.12 | 135 |
| 3-Hydroxyvaleric acid | 0.16 | 136 |
| 3-Phenylthiopropanol | 0.08 | 137 |
| Aniline | 0.22 | 138 |
| 4-Hydroxy-3-methoxyphenylglycol | 0.08 | 139 |
| 2-Methyl-2(p-methoxy)mandelate | 0.18 | 140 |
| 3,4-Dihydroxybenzeneacetic acid | 1.09 | 141 |
| 1-(2'-Hydroxyphenyl)-1-pentanol | 0.17 | 142 |
| Ethyl 3,4-dihydroxymandelate | 0.17 | 143 |
| 2,4-Dihydroxybenzoic acid | 0.18 | 144 |
| 3-Hydroxy-3-(4'-hydroxy-3'-methoxyphenyl)propionic acid | 0.28 | 145 |

| | | |
|---|---|---|
| Dodecanedioic acid | 0.13 | 146 |
| Silychristin A | 0.28 | 147 |
| Isolariciresinol | 0.31 | 148 |
| 3,3-dimethyl-4-methylene-1,2-bis(hydroxymethyl) Cyclopentene | 0.16 | 149 |
| Dihydroferulic acid | 0.46 | 150 |
| o-Methoxymandelic acid | 0.22 | 151 |
| 3,3-dimethyl-4-methylene-1,2-bis(hydroxymethyl) Cyclopentene | 0.16 | 152 |
| 3-Vanil-1,2-dihydroxypropane | 0.44 | 153 |
| Mandelic acid | 0.22 | 154 |
| Ethyl 3-(4-hydroxy-3-methoxyphenyl)lactate | 0.08 | 155 |
| 4-Hydroxy-3-methoxyphenylglycol | 0.09 | 156 |
| 4-Hydroxy-3-methoxyphenylglycol | 0.11 | 157 |
| Aniline | 0.14 | 158 |
| trans-Coniferyl alcohol | 0.08 | 159 |
| N-Phenylglycine | 0.19 | 160 |
| 4-hydroxy-3-methoxyphenethylamine | 1.44 | 161 |
| S-n-butylthiobenzoic acid | 0.25 | 162 |
| 4-(Hexyloxy)aniline | 0.2 | 163 |
| Aniline | 0.32 | 164 |
| 5-pentyl-3-hydroxyimino-1H-Indole-2,3-dione | 0.28 | 165 |
| 3,3-dimethyl-4-methylene-1,2-bis(hydroxymethyl) Cyclopentene | 0.11 | 166 |
| Aniline | 0.1 | 167 |

| | | |
|---|---|---|
| Aniline | 0.12 | 168 |
| 3-methoxy-α,4-dihydroxybenzenepropanoic acid | 0.16 | 169 |
| 3-methoxy-α,4-dihydroxybenzenepropanoic acid | 0.17 | 170 |

Γ - The peak number assigned to GC-MS chromatogram peaks in Figure 1, * - similarity percentage based on NIST (National Institute of Standards and Technology) 2017 GC-MS mass spectral library were used for spectral match.

**Table S4:** SSH composition (g per 100 g Dry Biomass$^{\Delta}$ (DW)).

| SSH Profile | | % DW |
|---|---|---|
| Total sugars | Arabinose | 19.3±0.7 |
| | Rhamnose | 1.0±0.0 |
| | Galactose | 10.6±0.4 |
| | Glucose | 5.6±0.2 |
| | Xylose | 27.3±1.2 |
| | Mannose | 23.1±1.1 |
| Inhibitors | Acetic Acid | 2.4±0.0 |
| | Formic Acid | 0.1±0.0 |
| | Levulinic acid | 0.2±0.0 |
| | Furfurals | 0.01±0.0 |
| | 5-Hydroxymethylfurfural | 0.2±0.0 |
| Total phenolics | LDMOs and extractives | 9.3±0.0 |
| Salts | Metal ions | 0.2±0.0 |

| | |
|---|---|
| **Total** | 99.4±1.8 |

Δ - Volatile compounds acetic acid, formic acid, furfural, HMF are not actually present in the lyophilized solids but are included in the normalized composition of dry biomass.
